# Weaker neural suppression in autism

**DOI:** 10.1101/645846

**Authors:** Michael-Paul Schallmo, Tamar Kolodny, Alexander M. Kale, Rachel Millin, Anastasia V. Flevaris, Richard A.E. Edden, Jennifer Gerdts, Raphael A. Bernier, Scott O. Murray

**Affiliations:** Department of Psychology, University of Washington, Seattle, WA; Department of Psychiatry and Behavioral Science, University of Minnesota, Minneapolis, MN; Department of Radiology and Radiological Science, Johns Hopkins University, Baltimore, MD; Department of Psychiatry and Behavioral Sciences, University of Washington, Seattle, WA

## Abstract

Increased neural excitation resulting from weakened inhibition is a leading hypothesis for the pathophysiology of autism. However, experimental support in humans remains equivocal. Alternatively, modulatory processes that suppress neural responses but do not specifically rely on inhibition may be impacted in ASD. Leveraging well-characterized suppressive neural circuits in the visual system, we used behavioral and fMRI tasks to demonstrate a significant reduction in neural suppression in young adults with ASD compared to neurotypical controls. We further tested the mechanism of this suppression by measuring levels of the inhibitory neurotransmitter GABA, and found no differences in GABA between groups. We show how a computational model that incorporates divisive normalization, as well as narrower top-down gain (that could result, for example, from a narrower window of attention), can explain our observations and divergent previous findings. Thus, weaker neural suppression in ASD may be attributable to differences in top-down processing, but not to differences in GABA levels.

## Introduction

Autism spectrum disorder (ASD) is a neurodevelopmental condition characterized by social difficulties, repetitive behaviors, and sensory abnormalities^5^, the cause of which remains unknown. Prominent theories suggest that ASD may result from a pervasive reduction of neural inhibition^6-8^, leading to an increase in the ratio of excitation/inhibition (E/I) throughout cortex. Reduced inhibition might thus account for symptoms including sensory over-responsivity in ASD. However, evidence for a widespread reduction in inhibition is either indirect or inconclusive. For example, there is evidence for decreased inhibition from genetic models of ASD in mice^7, 9-11^, but results in humans remain equivocal^8, 12-17^. An alternative is that modulatory processes that suppress neural responses, but do not rely directly on neural inhibition, are either disrupted or differentially engaged in ASD. This alternative account predicts that weaker suppression in ASD may not be widespread and would manifest only under specific circumstances. For example, in the visual system, suppressive modulatory mechanisms are engaged when a stimulus extends beyond the spatial receptive field of a given neuron^18, 19^. Although it has been suggested that suppressive visual surround modulation reflects neural inhibition^20^, work from our group and others has recently called this assumption into question^1, 21, 22^. Importantly, these suppressive effects can be modeled within the context of divisive normalization^23-25^, a computational framework for characterizing modulatory processes in visual cortex.

In the current study, we examined visual spatial suppression in ASD at both a behavioral and neural level, using visual psychophysics and functional MRI (fMRI) respectively. Additionally, we used MR spectroscopy to measure neurotransmitter levels *in vivo*, in order to probe inhibition. Our results indicate that ASD is associated with abnormally weak neural suppression within the motion-sensitive brain area called human MT complex^26, 27^ (hMT+), but that this reduction in neural suppression is not driven by abnormal GABA levels in the hMT+ region. A model that incorporates divisive normalization and narrower top-down gain provides a parsimonious computational basis for the observed differences in suppression in ASD.

## Results

To assess suppressive modulatory mechanisms in humans with ASD, we measured visual spatial suppression, a phenomenon in which larger moving stimuli are more difficult to perceive^28^. This mirrors a well-known neural phenomenon; when stimuli extend beyond a neuron’s spatial receptive field, neural responses in visual cortex are suppressed through a combination of feed-forward, lateral, and feedback interactions^18, 19, 29^. Based on work in both humans^1, 30, 31^ and non-human primates^32-35^, it is thought that neural surround suppression within the motion-selective visual area MT plays an important role in the perceptual phenomenon of spatial suppression during motion discrimination. In a series of 3 experiments within the current study, we characterized spatial suppression in terms of behavioral, neural, and neurochemical differences between a group of 28 young adults with ASD and a comparison group of 35 neurotypical (NT) participants (for demographic information, see Table 1). We then present a computational account of weaker spatial suppression in the context of a divisive normalization model.

**Table 1.** Participant demographics. Values shown are group mean and *SD*. Results of statistical tests for group differences are shown on the right.

| Demographics | ASD (n = 28) | NT (n = 35) | Statistics |
| --- | --- | --- | --- |
| Age in years | 22.4 (3.44) | 23.4 (3.56) | $t_{(61)} = 1.09, p = 0.3$ |
| Biological sex | 18 M; 10 F | 21 M; 14 F | $\chi^2_{(1)} = 0.008, p = 0.9$ |
| Non-verbal IQ<sup>†</sup> | 112 (17.6) | 113 (13.2) | $t_{(61)} = 0.418, p = 0.7$ |
| Handedness | 4 L; 24 R | 2 L; 33 R | $\chi^2_{(1)} = 0.518, p = 0.5$ |
| ADOS-2 score<sup>‡</sup> | 7.32 (1.59) | - | - |

<sup>†</sup>Non-verbal IQ was calculated based on the Wechsler Abbreviated Scale of Intelligence (WASI).
<sup>‡</sup>Autism Diagnostic Observation Schedule, 2<sup>nd</sup> Ed. (ADOS-2) comparison score.

### Behavior

We obtained a quantitative behavioral index of spatial suppression by measuring motion duration thresholds^28^. It is known that the amount of time that a stimulus needs to be presented in order to perceive motion direction depends on stimulus size; paradoxically, larger stimuli require longer presentation durations^36^. In our task, participants judged whether visual grating stimuli drifted left or right (Figure 1A-C). Motion duration thresholds were defined by the minimum stimulus duration for which participants could perceive motion direction with 80% accuracy. Thresholds were measured for each of 3 different stimulus sizes and 2 different contrasts (Figure 1A & B). Note that this task does not depend on reaction time; although stimulus duration was brief (Figure 1C), response time was not limited.

**Figure 1.**
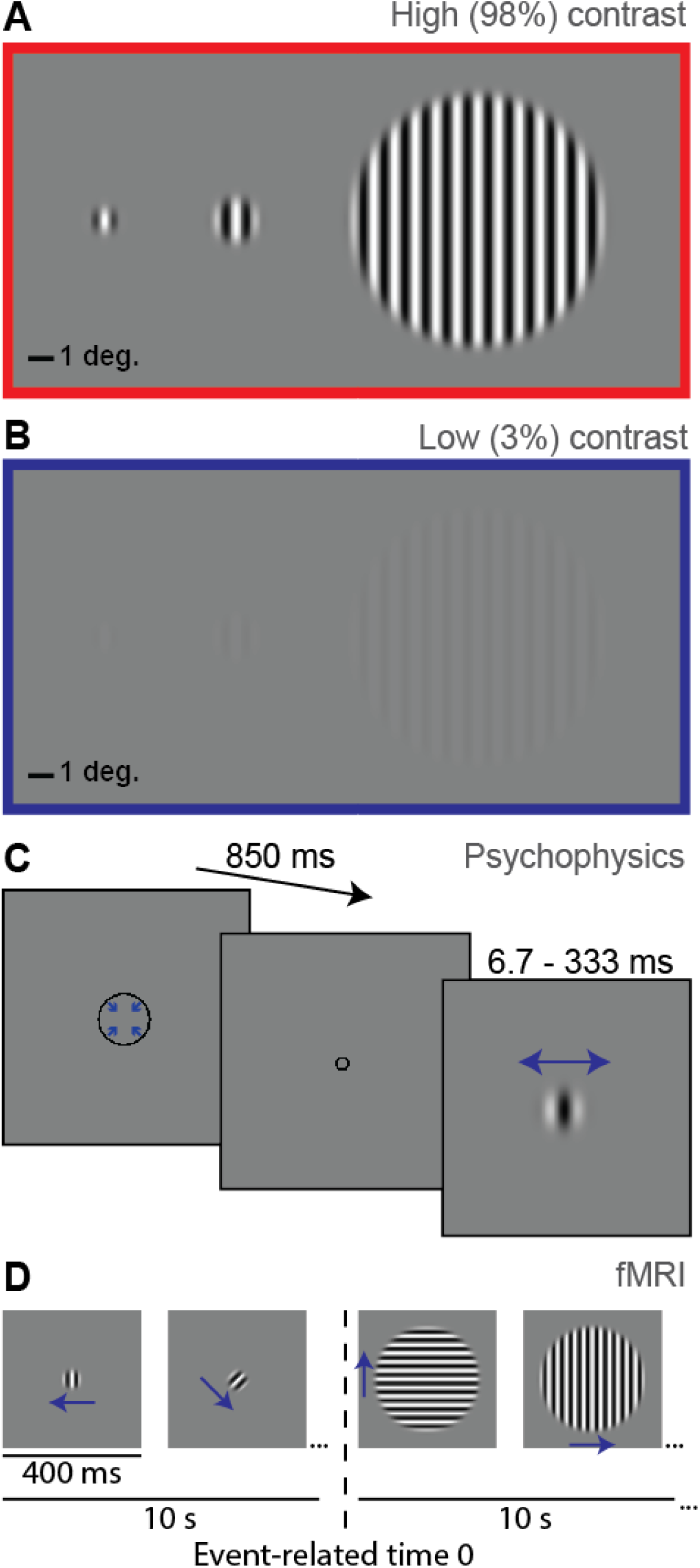
Stimuli and paradigms. **A)** small (0.84°), medium (1.7°) and big gratings (10° diameter) at high contrast (98%). **B)** the same gratings at low contrast (3%). **C)** Psychophysical paradigm; a fixation mark (shrinking circle) was followed by a briefly presented drifting grating (left or right; small high contrast grating shown). Blue arrows indicate direction of motion. **D)** FMRI paradigm; alternating 10 s blocks of smaller (2°) and larger (12°) gratings (high contrast shown). Vertical dashed line indicates the transition from smaller to larger, which is the event of experimental interest. Note that stimuli are scaled differently across panels for display purposes.

We observed the expected spatial suppression effect; thresholds were significantly longer for larger stimuli (main effect of size; *F*_1, 61_ = 44.1, *p* = 9 × 10^-9^; Figure 2A & B). Importantly, spatial suppression was significantly weaker in the ASD group vs. NTs (group x size interaction; *F*_1, 61_ = 9.76, *p* = 0.003). Thus, as stimulus size increased, duration thresholds increased less dramatically among participants with ASD. Weaker spatial suppression in ASD did not depend on stimulus contrast; other interactions and main effects, including a main effect of group, were also not significant (all *F*_1,61_ < 1.78, *p*-values > 0.19).

**Figure 2.**
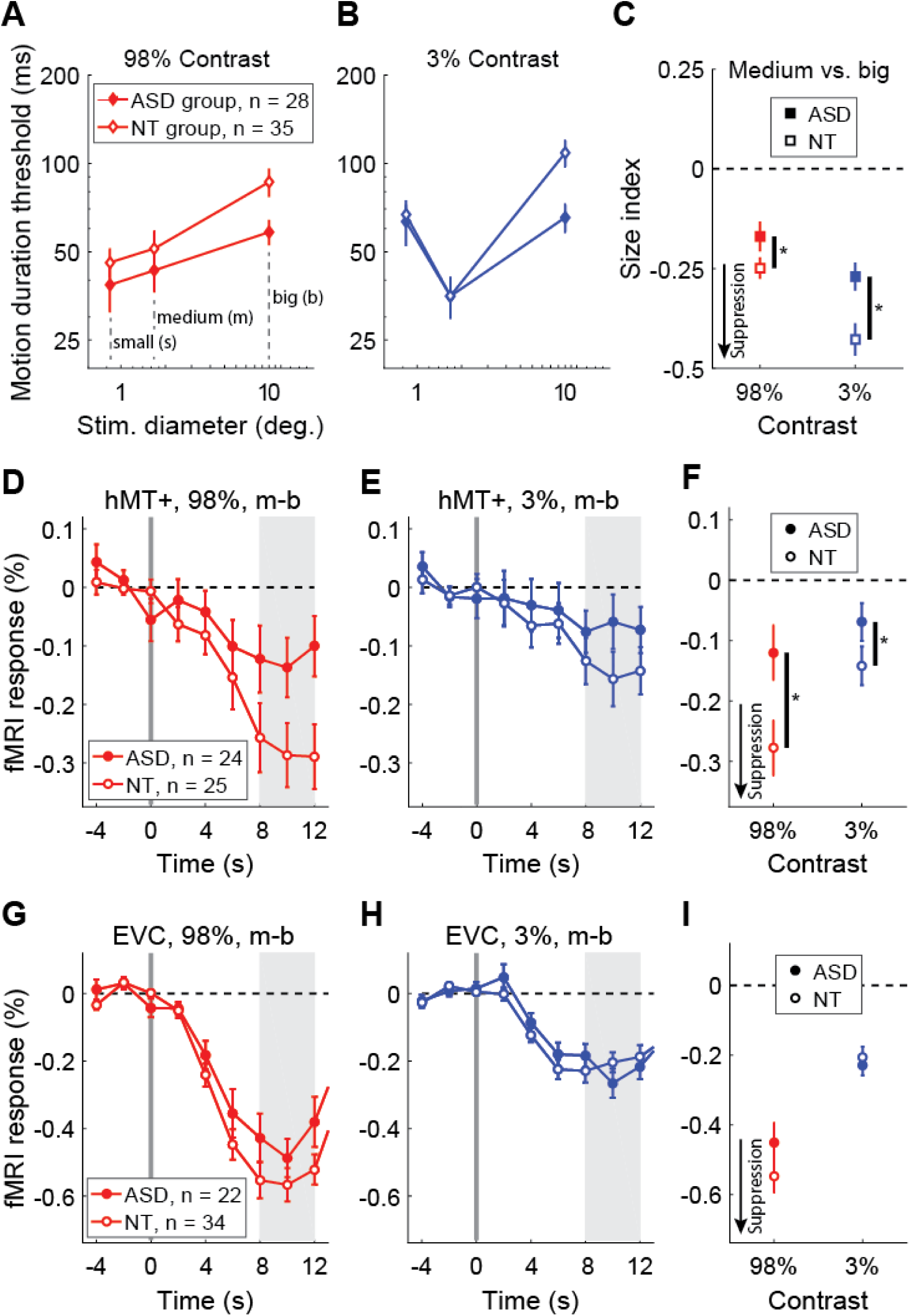
Behavioral and fMRI results. **A)** Motion duration thresholds for high contrast (98%) drifting gratings. The time required to perceive whether stimuli drifted left or right with 80% accuracy is shown on the y-axis for different stimulus sizes (x-axis). **B)** The same, but for low contrast gratings (3%). **C)** The effect of increasing stimulus size on duration thresholds was quantified using a size index – the difference between log thresholds (Equation 3). Negative values indicate suppression. Participants with ASD show weaker spatial suppression during motion discrimination than NTs. **D)** FMRI responses in foveal human MT complex (hMT+) to an increase in stimulus size; at time = 0 s, high contrast drifting gratings increased in size from medium (m) to big (b). **E)** The same, but for low contrast gratings. **F)** Average fMRI responses for each group (from shaded regions in **E** & **F**). Panels **G**-**H** are the same as **D**-**F**, but for a foveal region of early visual cortex (EVC). Samples sizes for each row are shown in **A, D**, & **G**. Error bars are *S.E.M.*

To quantify spatial suppression we computed size indices, which involved taking the logarithm and then the difference between thresholds for medium and large stimuli (see Equation 3 in the Methods). More-negative size indices reflect stronger suppression (greater increase in duration thresholds with increasing stimulus size; Figure 2C). Comparing size indices between groups showed the same effect of weaker spatial suppression in participants with ASD vs. NTs (main effect of group; *F*_1, 61_ = 9.37, *p* = 0.003). In this case, we found size indices in both groups were more negative for low vs. high contrast gratings, indicating stronger suppression (main effect of contrast; *F*_1, 61_ = 19.7, *p* = 4 × 10^-5^). Weaker suppression in ASD did not depend on stimulus contrast (group x contrast interaction; *F*_1, 61_ = 1.55, *p* = 0.2). Together, these results indicate that our participants with ASD experienced weaker spatial suppression; they were able to perceive the direction of motion for stimuli presented more-briefly, especially when those stimuli were large (the most challenging condition). Weaker spatial suppression in our behavioral task suggests that neural suppression in visual cortex may also be weaker in ASD.

### Functional MRI

We used an fMRI paradigm designed to measure spatial suppression in order to probe neural suppression in ASD more directly. In NT subjects, we have recently shown that this paradigm, which involves presenting alternating blocks of smaller and larger drifting gratings (Figure 1D), yields suppressed fMRI responses for larger vs. smaller stimuli within foveal regions of visual cortex^1^. Participants performed a colored-shape detection task at fixation, in order to minimize eye movements, divert attention away from the drifting gratings, and emphasize bottom-up stimulus processing.

We first examined the fMRI response within the motion-selective region of the lateral occipital lobe known as human MT complex (hMT+; Figure 3A & B). Because hMT+ is retinotopically organized^37^, and because our stimuli were presented at the fovea, we focused on voxels that showed significant selectivity for foveal over peripheral stimuli (see Methods; Supplemental Figure 1). For both low and high contrast stimuli, fMRI responses in foveal hMT+ to larger stimuli were significantly below baseline (i.e., lower than responses to the preceding smaller stimuli; paired *t*-tests, *t*_47_ > 4.13, *p*-values < 0.003, Bonferroni corrected for 2 comparisons; Figure 2D & E). Thus, we saw robust suppression of the fMRI signal in hMT+ in response to increasing stimulus size, in agreement with the spatial suppression effect observed in our motion discrimination task above. Importantly, fMRI suppression within hMT+ was significantly weaker for ASD vs. NT participants (main effect of group, *F*_1, 47_ = 5.66, *p* = 0.022; Figure 2F). Suppression in both groups was stronger for high vs. low contrast stimuli (main effect of contrast, *F*_1, 47_ = 5.52, *p* = 0.023), but weaker suppression in ASD did not depend on contrast (group x contrast interaction, *F*_1, 47_ = 1.15, *p* = 0.3). These findings indicate that neural suppression in foveal hMT+ is weaker for participants with ASD compared to their NT peers, in agreement with our behavioral results.

**Figure 3.**
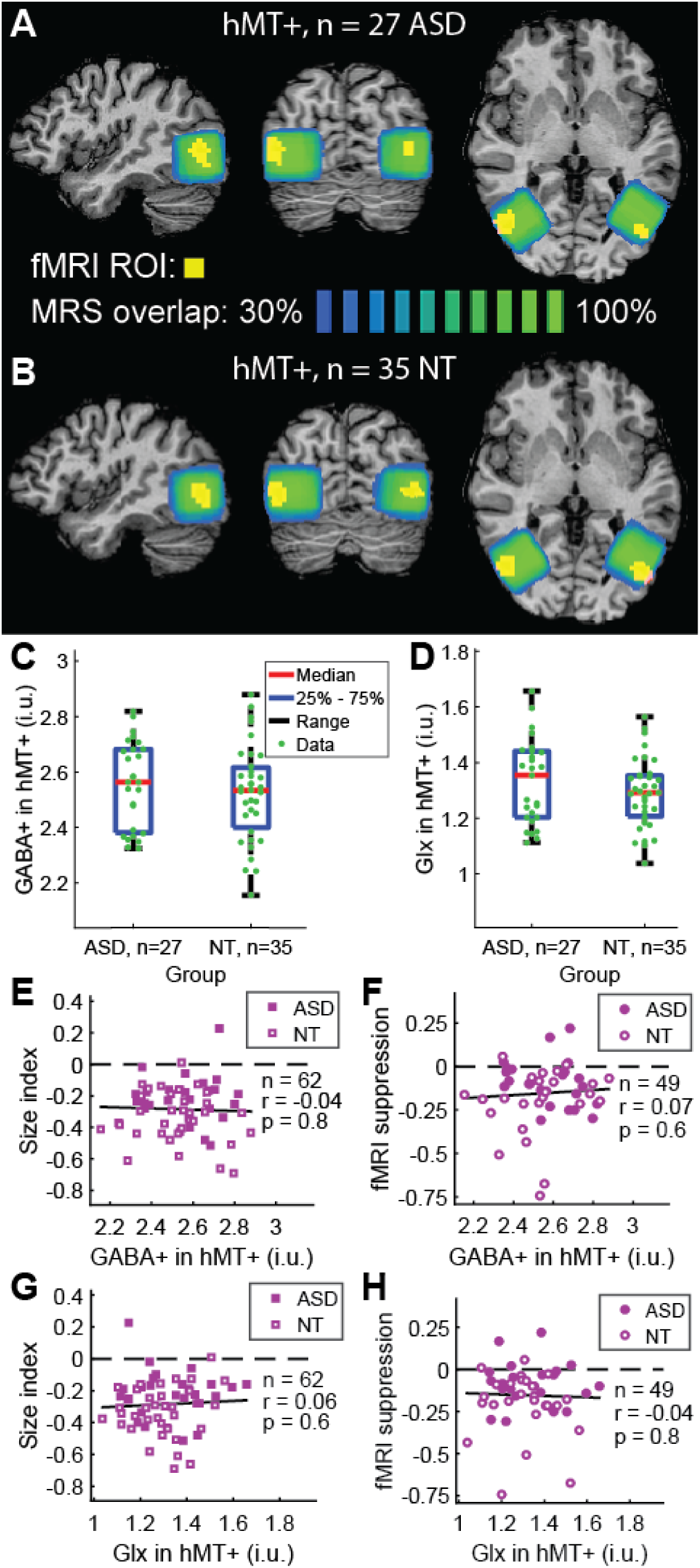
MR spectroscopy results. **A)** Voxel positions for left and right hMT+ in Talairach space for n = 27 participants with ASD. Blue-green color indicates overlap across individuals. Yellow indicates average hMT+ ROI across participants, from fMRI (correlation of predicted vs. observed response, visualization thresholds: *r*123 ≥ 0.25). **B)** Same, but for n = 35 NTs. **C)** GABA+ concentrations in hMT+ (averaged left and right voxels in each participant) in institutional units (i.u.). **D)** Same, but for Glx. **E & F)** Correlation between GABA+ in hMT+ and size indices or fMRI suppression (averaged across low & high contrast), respectively. **G & H)** Same, but for Glx.

Next, we examined fMRI responses within a region of early visual cortex (EVC; at the foveal confluence of V1, V2, and V3 near the occipital pole). We have previously found that among NT individuals, fMRI responses in foveal EVC are also suppressed by larger vs. smaller stimuli, but that the pattern of fMRI suppression in hMT+ was a better match to the spatial suppression observed psychophysically^1^. Here again, fMRI responses in EVC were significantly suppressed below baseline for both high and low contrast stimuli (paired *t*-tests, *t*_54_ > 9.04, *p*-values < 4 × 10^-11^, Bonferroni corrected for 2 comparisons; Figure 2G & H), in agreement with the expected spatial suppression effect. However, unlike in hMT+, we found no significant difference in fMRI suppression within EVC between participants with ASD and NTs (main effect of group, *F*_1, 55_ = 1.61, *p* = 0.2; Figure 2I). Suppression in EVC was stronger for high vs. low contrast stimuli (main effect of contrast, *F*_1, 55_ = 56.5, *p* = 7 × 10^-10^), but there was no significant interaction between group and contrast (group x contrast interaction, *F*_1, 53_ = 1.50, *p* = 0.2). These results indicate that fMRI responses within EVC reflect spatial suppression, but unlike for motion discrimination or responses in hMT+, there was no difference in fMRI suppression within EVC between participants with ASD and NTs.

### MR spectroscopy

After observing weaker neural suppression in the foveal hMT+ fMRI response in participants with ASD, we asked whether this difference in suppression might be attributable to weaker inhibition, consistent with theories of E/I imbalance in this disorder^6-8^. We used MR spectroscopy to measure the concentration of GABA+ (GABA, an inhibitory neurotransmitter, plus co-edited macromolecules) in a region centered around hMT+ in the lateral occipital lobe (Figure 3A & B; Supplemental Figure 5). We predicted that if weaker spatial suppression was driven by reduced inhibition in hMT+, this might be reflected in lower GABA+ levels as measured by MRS. However, we found no significant difference in GABA+ levels within the MT region between participants with ASD versus NTs (main effect of group, *F*_1, 60_ = 0.64, *p* = 0.4; Figure 3C). We also found no significant correlations between GABA+ levels in hMT+ and measures of suppression (size indices or fMRI; averaging across low & high contrast conditions to minimize multiple comparisons; correlations for both groups combined: |*r*_47-60_| < 0.08, uncorrected *p*-values > 0.6; for ASD participants alone: |*r*_22-25_| < 0.27, uncorrected *p*-values > 0.19; for NTs alone: |*r*_23-33_| < 0.19, uncorrected *p*-values > 0.3; Figure 3E & F). This agrees with our recent findings from a subset of these NTs^1^. These results suggest that weaker spatial suppression in ASD may not be attributed to differences in the level of GABA+ in hMT+, as measured by MRS.

Our MRS procedure also allowed us to measure a signal associated with glutamate, an excitatory neurotransmitter. Because this signal also includes some contribution from glutamine and glutathione^38^, we refer to this measure as Glx. Some previous studies have suggested that surround suppression is driven primarily by a withdrawal of excitation, rather than inhibition of neural activity^21, 39^. Therefore, we next examined whether weaker suppression in ASD might be driven by differences in excitatory neurotransmitter levels, which could be reflected in the Glx signal from the hMT+ region. However, we observed no significant difference between groups in Glx within hMT+ (main effect of group; *F*_1, 60_ = 1.22, *p* = 0.3; Figure 3D). Further, Glx levels in hMT+ did not correlate significantly with behavioral or fMRI measures of suppression (correlations for both groups combined: |*r*_47-60_| < 0.06, uncorrected *p*-values > 0.6; ASD participants alone: |*r*_22-25_| < 0.10, uncorrected *p*-values > 0.6; NTs alone: | *r*_23-33_| < 0.13, uncorrected *p*-values > 0.5; Figure 3G & H). These results do not lend support to the notion that differences in glutamate levels in hMT+, as indexed by Glx from MRS, underlie weaker spatial suppression in ASD.

Additional analyses revealed no significant interactions between participant group and the hemisphere (left or right hMT+) in which MRS data were acquired (see Supplemental information for full details). MRS signals (GABA + & Glx) were quantified relative to water (see Methods). Using water as a reference for scaling did not qualitatively affect our results (see Supplemental information). We also found no significant group differences in GABA+ or Glx within a second occipital MRS voxel location (medial region of early visual cortex; Supplemental Figure 2), and no significant correlations between suppression measures and levels of GABA+ or Glx in EVC (see Supplemental information).

### Relation to clinical measures

To probe whether weaker spatial suppression in ASD is related to clinical functioning, we first examined correlations between the total comparison score on the Autism Diagnostic Observation Schedule, 2^nd^ Edition (ADOS-2)^40^ and behavioral or fMRI measures of suppression. No significant correlations between ADOS-2 total comparison scores and either size indices or fMRI suppression in hMT+ were observed in our participants with ASD (|*r*_22-26_| < 0.18, uncorrected *p*-values > 0.3).

Abnormal sensory experiences are common among people with ASD^5^. Therefore, we also examined whether aspects of sensory processing in everyday life, as measured by the Sensory Profile^41, 42^ sub-scales for sensitivity and avoiding (summed scores), were associated with suppression metrics in both ASD and NT participants. These measures reflect ranked self-reported responses to questions such as “I am bothered by unsteady or fast-moving images.” We observed a moderate correlation between higher sensory sensitivity + avoiding and weaker fMRI suppression in hMT+ (*r*_46_ = 0.34, uncorrected *p* = 0.019, Bonferroni corrected for 4 multiple comparisons between suppression metrics and symptom scores *p* = 0.074; Figure 4). However, no significant relationship with behavioral size indices was found (*r*_60_ = -0.004, uncorrected *p* = 0.98). The former result may suggest that weaker neural suppression within hMT+ could be relevant to sensory dysfunction during daily life.

**Figure 4.**
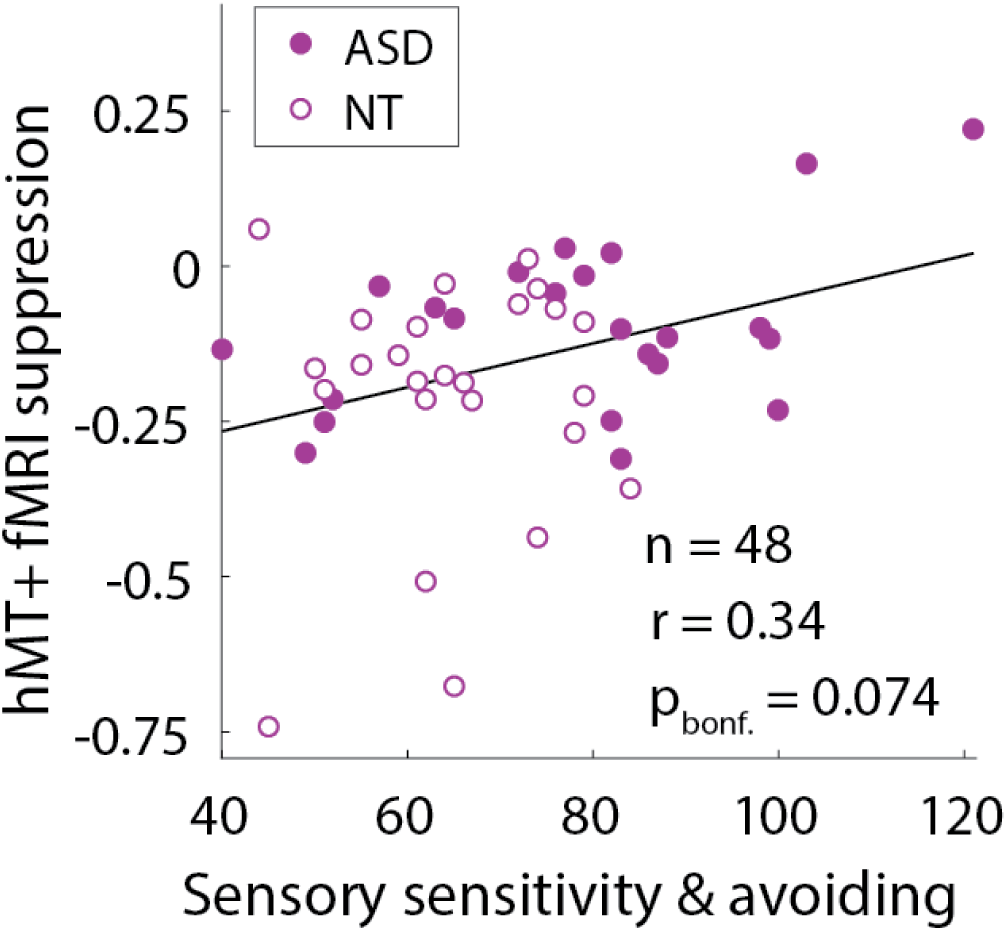
Relationship with sensory dysfunction. Weaker fMRI suppression in foveal hMT+ was associated with higher combined scores for sensory sensitivity and avoiding from the Sensory Profile, across both ASD and NT participants.

### Computational modeling

We next examined different computational principles that might account for weaker neural suppression in ASD. Recent work^1, 2^ has suggested that a general computational model for spatial vision, known as divisive normalization (Figure 5), can describe the effect of stimulus size on motion duration thresholds measured psychophysically. Divisive normalization models have been used to describe the effects of spatial context on neural responses in visual cortex, which can be summarized by saying: a neuron’s response is divided by the summed response of its neighbors^23, 24^. Rosenberg and colleagues^2^ have proposed that a reduction in divisive normalization might account for abnormal performance across a number of visual tasks in people with ASD. Weaker normalization in ASD would be expected to yield superior motion discrimination performance (i.e., lower duration thresholds), as previously reported^3^. Another computational model has been presented by Schauder and colleagues^4^, who suggested that larger excitatory spatial filters could account for their observation of higher motion duration thresholds among young people with ASD vs. NTs. We have recently used a divisive normalization model to describe spatial suppression across a series of experiments in NT participants^1^. Here, we considered three different versions of this model, including variants based on the two models noted above^2, 4^, in an attempt to describe our current observations of weaker spatial suppression in ASD within a normalization model framework.

**Figure 5.**
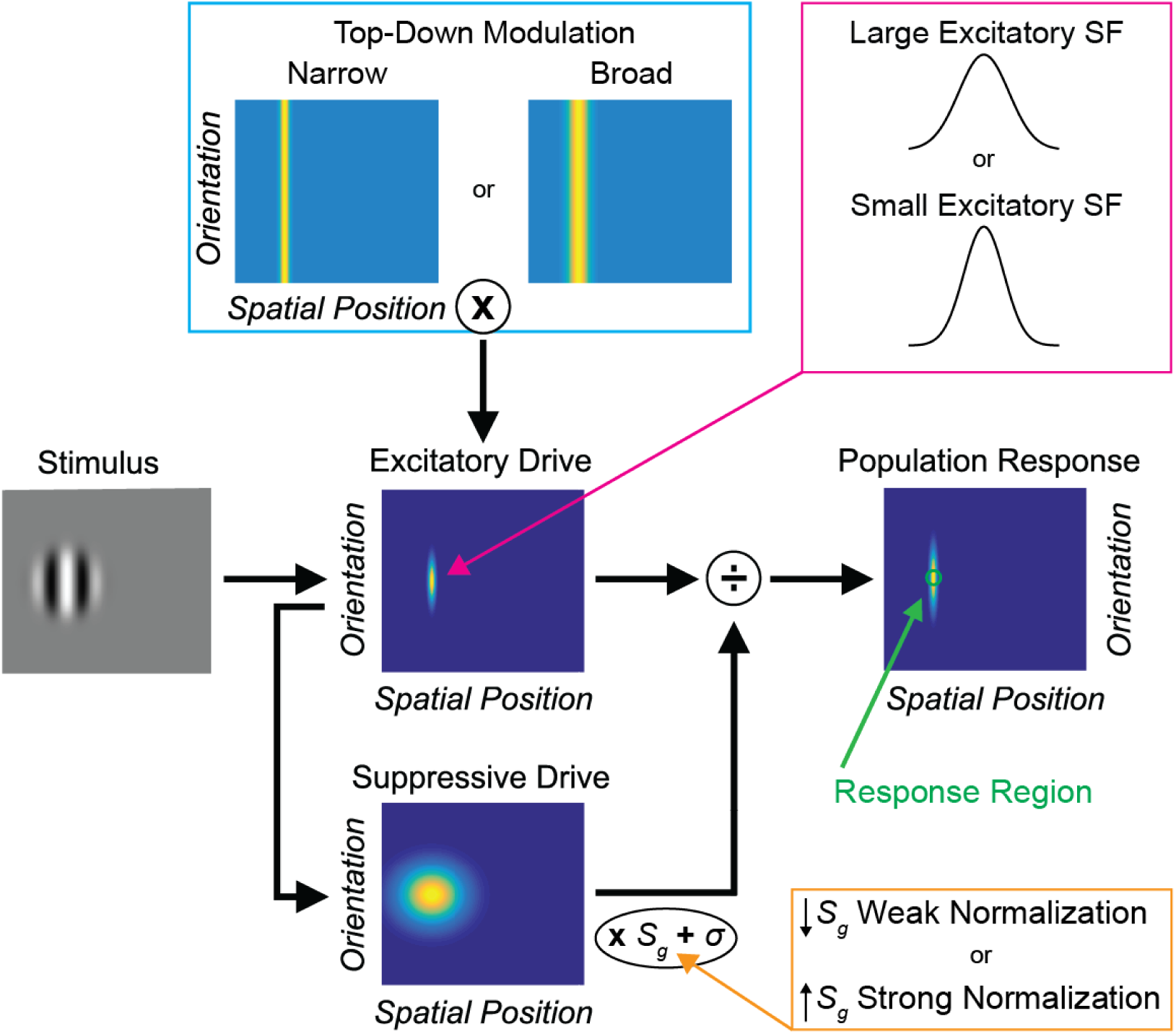
Normalization model diagram. For full model details, see Methods and Supplemental information. We tested three different model variants, which are described within colored boxes. **Orange**: Weaker vs. stronger normalization was modeled by lower vs. higher values for the suppressive gain term (*Sg*), as in previous work^2^. **Magenta**: Large vs. small excitatory spatial filters (SFs) were modeled using wider vs. narrower spatial Gaussians in the excitatory drive term (*E*), as in previous work^4^. **Cyan**: Narrow vs. broad top-down modulation was modeled using narrower vs. broader spatial Gaussians in the top-down modulation term (*M*).

We first asked whether weaker normalization strength (i.e., a 25% reduction in suppressive gain^2^; Figure 5, orange box; Supplemental Table 2) could describe the difference in spatial suppression we observed psychophysically between individuals with ASD and NTs. We found that although weaker normalization reduced the motion duration thresholds predicted by the model (Figure 6A & B), this reduction (observed across all stimulus sizes) was not specific to the largest stimulus. Thus, weaker normalization did not dramatically reduce spatial suppression (Figure 6C), unlike our psychophysical results in participants with ASD (Figure 2A-C). Therefore, we conclude that a reduction in suppressive gain within the normalization model^2^ is not sufficient to account for our observations of weaker spatial suppression in participants with ASD.

**Figure 6.**
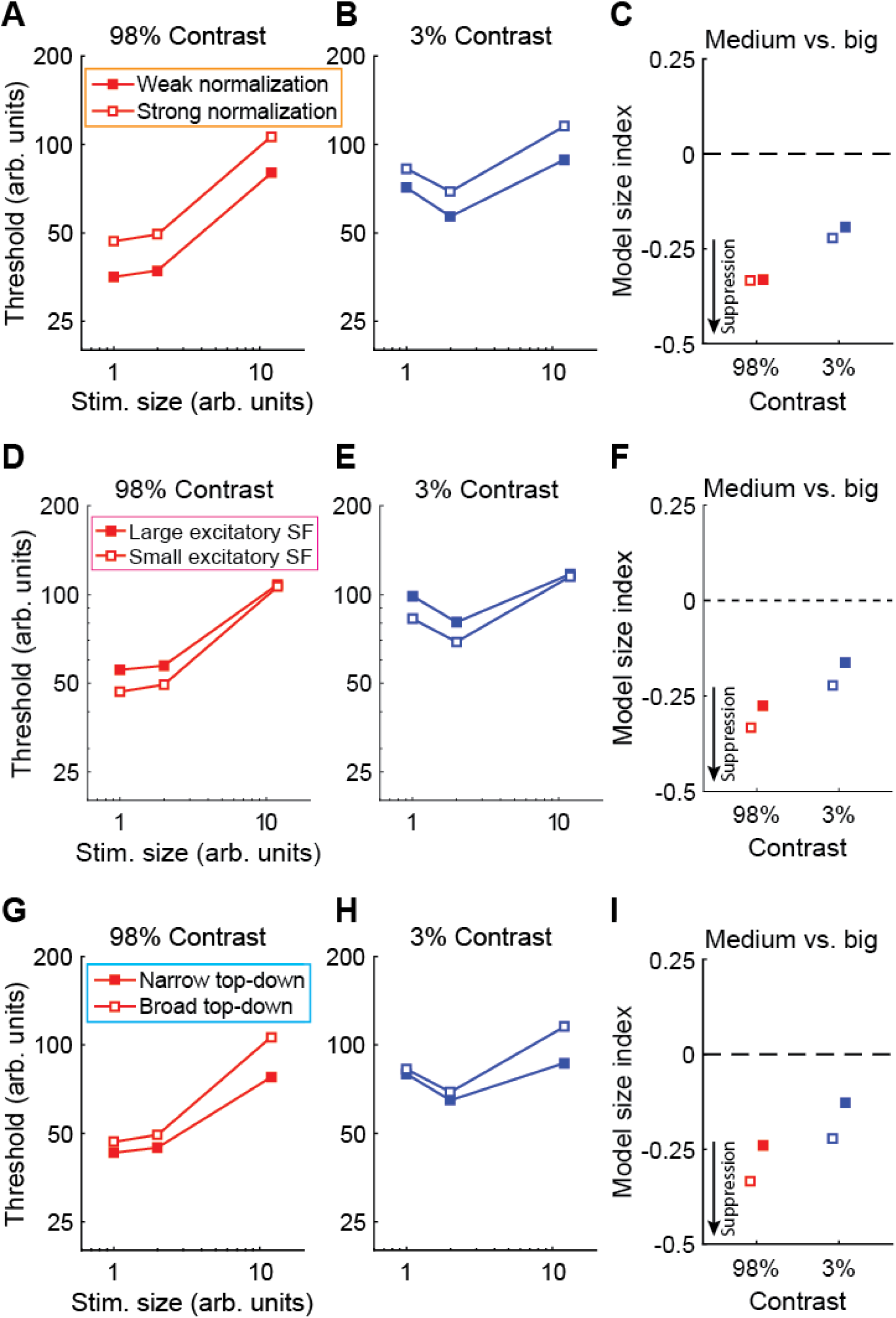
Normalization model results. We used a divisive normalization model to describe motion duration thresholds, as in our previous work^1^. **A & B)** Weaker normalization (25% weaker suppressive gain^2^) yields lower predicted duration thresholds across stimulus sizes and contrasts. **C)** Size indices are not dramatically altered by weaker normalization. **D-F)** Larger excitatory spatial filters (25% larger^4^) yield higher duration thresholds and less-negative size indices. Thus, neither weak normalization nor larger excitatory spatial filters provide a good match for our observations of lower motion discrimination thresholds for large stimuli among people with ASD (Figure 2A-C). **G & H)** Narrower top-down modulation (6 vs. 14 arb. units) yields lower thresholds at larger stimulus sizes. **I)** Narrower top-down modulation produces less-negative size indices, consistent with weaker suppression, and thus shows a better match to our psychophysical results.

Next, we examined whether larger excitatory spatial filters (25% wider^4^; Figure 5, magenta box; Supplemental Table 2) could account for weaker spatial suppression within our model framework. Indeed, we found that larger excitatory spatial filters did predict weaker spatial suppression (Figure 6F). However, this was driven by larger predicted duration thresholds for small and medium sized stimuli (Figure 6D & E), unlike the pattern of results we observed in participants with ASD (smaller thresholds for large stimuli; Figure 2A & B). Therefore, we find that larger excitatory spatial filters^4^ within the framework of our normalization model are not sufficient to explain the pattern of motion duration threshold data we observed in people with ASD.

Finally, we considered whether differences in top-down gain modulation (that could, for example, reflect differences in spatial attention), as described by the normalization model, might better account for our observations of weaker spatial suppression in ASD (Figure 2A-C). It has been suggested that the focus of spatial attention may be narrower in people with ASD^43^, and top-down effects (such as attention or expectation) that modulate the gain of neural processing can be modeled in terms of divisive normalization^24^. We found that a narrower top-down gain field within our model (6 vs. 14 arbitrary units; Figure 5, cyan box; Supplemental Table 2) led to predicted motion duration thresholds that were smaller, especially for larger stimuli (Figure 6G & H). Model size indices were likewise less-negative with narrower top-down modulation (Figure 6I), indicating weaker spatial suppression. Narrower top-down gain modulation within the normalization model therefore predicts a pattern of results that closely mirrored our observation of weaker spatial suppression in ASD (Figure 2A-C). We also found that using even narrower top-down parameters (1 or 2 vs. 6 arbitrary units) yielded model predictions that showed similarities to previous observations of both lower^3^ or higher^4^ motion duration thresholds in people with ASD vs. NTs (see Supplemental information; Supplemental Figure 3). Thus, narrower spatial top-down gain modulation within the normalization model framework may be able account for our observation of weaker spatial suppression among participants with ASD vs. NTs, as well as previous divergent findings.

### Control analyses

Finally, we performed a series of control analyses to rule out alternative explanations for our results showing weaker behavioral and fMRI suppression in the ASD group (Figure 2A-F). Although demographic factors did not differ significantly between groups, previous work has shown that age^44-48^, biological sex^49^, and IQ^49-51^ may each be associated with differences in motion duration thresholds and / or the magnitude of spatial suppression. We sought to control for these factors in post-hoc analyses by including them as covariates when testing for group differences in behavioral or fMRI suppression. Our results were unaffected by including age, sex, and IQ as factors in linear mixed effects models; suppression was still significantly weaker among participants with ASD vs. NTs in both the motion discrimination task and in the fMRI response within hMT+ (see Supplemental information).

We also asked whether our fMRI results might be explained by differences in head motion or fixation task performance between groups. To address this, we excluded fMRI data with excessive head motion or poor fixation task performance (see Supplemental information). However, the results were qualitatively the same as those shown in Figure 2G-I; after exclusion, fMRI response suppression within hMT+ was weaker among participants with ASD vs. NTs (Supplemental Figure 4).

Last, we examined whether differences in eye movements between groups might explain weaker spatial suppression in participants with ASD. We found no significant differences in eye movement metrics between groups, and no correlations between these metrics and our measures of suppression from psychophysics or from fMRI (see Supplemental information for results and methodological details). Together, the results of these control analyses do not support these alternative explanations of our observations of weaker spatial suppression among participants with ASD.

## Discussion

We have found evidence for weaker neural suppression in people with autism. Specifically, spatial suppression, a phenomenon we measured in both a visual motion discrimination task and using functional MRI in the visual area hMT+, was significantly weaker among participants with ASD compared with demographically-matched NT individuals (Figure 2A-F). While it may be tempting to attribute weaker neural suppression in ASD to weaker inhibition, we found no evidence to support this notion, as GABA levels were equivalent among both groups (Figure 3C & D, Supplemental Figure 2C & D). Instead, our computational model suggests that weaker suppression in ASD may have an excitatory basis: top-down gain, which modulates excitation, may be tuned more narrowly in space in persons with ASD (Figure 5, cyan box; Figure 6G-I).

Reduced inhibition, leading to a net increase in the ratio of excitation/inhibition, is frequently hypothesized as a core pathophysiological mechanism in autism^6-8^. It has also been suggested that abnormal visual suppression observed in clinical populations may reflect dysregulation in the strength of underlying inhibitory mechanisms^16, 20, 44, 52-54^. However, we did not find evidence to suggest a difference in inhibition between participants with ASD and NTs; MR spectroscopy measurements of GABA+ in visual cortex did not differ between groups (Figure 3C and Supplemental Figure 2C), and were not correlated with neural or perceptual measures of suppression (Figure 3E-F). Nevertheless, we cannot rule out the possibility that the mixed nature of the MRS signal (i.e., GABA plus co-edited macromolecules) could have masked subtle underlying group differences in the neurotransmitter signals of interest. Given these caveats, we conclude that there is no evidence to support an overall group difference in GABA (or glutamate) in visual cortex as assessed by MRS in our hands, nor for any relationship between these metabolites and our fMRI or behavioral measures of suppressive modulation. Studies in animal models have shown that both excitatory and inhibitory neural mechanisms contribute to suppressive neural modulation in visual cortex^21, 22, 39, 55-58^. Consistent with recent pharmacological work in nonhuman primates^22^, our recent study using lorazepam (a GABA_A_ receptor modulator) in NT humans^1^ showed that strengthening inhibition yields *weaker*, rather than stronger spatial suppression during motion discrimination. Our current MRS results are in line with previous observations of normal GABA+ levels within visual cortex in people with autism^13, 15, 17^ (but see ^14^ for a more nuanced report). Thus, although there is evidence for weaker inhibition in mouse models of ASD^7, 9-11^, our results do not support the notion that weaker inhibition drives weaker neural suppression in humans with ASD.

Our computational modeling work suggests an alternative mechanism may explain weaker neural suppression in ASD: spatially narrower top-down gain modulation. The reason that narrower top-down gain can result in weaker neural suppression may be understood intuitively by referring to Figure 5 (cyan box). When stimuli are small, whether the top-down gain is narrow or broad has little effect on the model behavior; the suppressive drive, and the excitatory drive it depends on, are both similarly engaged. However, when a stimulus is large, broader top-down gain results in stronger engagement of the suppressive drive (which is spatially broad), yielding more-drastic increases in predicted thresholds for larger stimuli (i.e., stronger suppression). Likewise, narrower top-down gain predicts weaker suppression.

Narrower top-down neural gain could, for example, reflect intrinsic differences in spatial attention – individuals with autism may have narrower ‘windows-of-attention’ compared to NT individuals. While this is consistent with previous experimental findings showing a sharper gradient of spatial attention in ASD^43^ and is consistent with ‘detail-focused’ perceptual behavior associated with autism^59-61^, we did not explicitly manipulate spatial attention in the current study. Thus, we can only speculate about the cognitive origins of the narrower spatial gain suggested by the model. Importantly, the model does not imply any *impairment* or *reduction* in spatial attention in ASD *per se*, only a small difference in how attention is allocated in space. This difference in allocation could reflect an inclination towards local processing, as is sometimes invoked to characterize ASD^60^ or may reflect intrinsically altered structure of feedback circuits^18, 62^.

Weaker neural suppression in ASD does not seem to extend to earlier regions of visual cortex; we found weaker suppression among participants with ASD in the fMRI response within foveal hMT+ (Figure 2D-F), but not in the foveal region of early visual cortex (EVC) near the occipital pole (at the confluence of V1, V2, and V3; Figure 2G-I) that provides input to area MT. At first, this finding may appear at-odds with the notion of narrower top-down modulation in ASD, since top-down effects such as spatial attention are known to modulate responses in V1^63-68^. However, the magnitude of these modulatory effects varies greatly across different regions of visual cortex. In general, larger effects of top-down modulation have been observed in higher visual areas (like hMT+), versus smaller effects at earlier stages^69^. Thus, in the current study, top-down modulation might be reflected to a greater degree in the fMRI responses within hMT+, as compared to those in EVC. Future studies that investigate other specialized, later stages of processing (e.g., responses to face stimuli in fusiform cortex) will be better positioned to address the specificity of our findings, and determine (for example) whether weaker suppression in ASD is restricted to motion stimuli, or is a general feature of higher-level visual processing.

Weaker neural suppression in ASD could be expected to have important consequences for sensory processing in one’s daily life. A straightforward prediction is that reduced neural suppression would manifest in terms of increased sensory sensitivity. Our results bore this prediction out in a limited way; we found a modest correlation between higher sensory sensitivity + avoiding scores and weaker fMRI suppression within foveal hMT+ across both participant groups (Figure 4). Although this observation was limited to the visual system, it is known that abnormal sensory phenomena can occur across modalities in ASD^5^. Future research that examines neural suppression in different modalities (e.g., somatosensation and audition) may provide a clearer link between abnormal neural suppression and sensory symptoms.

## Methods

### Participants

Our study included 28 young adult participants on the autism spectrum (18 male, 10 female), and 35 neuro-typical comparison participants (21 male, 14 female). Data from these participants with ASD^70^ and NTs^1, 70, 71^ were included in our recently published work. Diagnoses were confirmed through the Autism Diagnostic Observation Schedule (ADOS)^40^, Autism Diagnostic Interview-Revised (ADI-R)^72^ and clinical judgment using DSM-5 criteria^73^. The following demographic factors did not differ significantly between the two participant groups: age, biological sex, non-verbal IQ (from the Wechsler Abbreviated Scale of Intelligence; WASI^74^), and handedness. See Table 1 for demographic information.

Our inclusion criteria were as follows: age 18-30 years, non-verbal IQ > 70, normal or corrected to normal visual acuity and no visual impairments, no impairment to sensory or motor functioning, no history of seizures or diagnosis of epilepsy, no neurological disease or history of serious head injury, no nicotine consumption in excess of 1 cigarette per day within the last 3 months, no use of illicit drugs within the past month, no consumption of alcohol within 3 days prior to MR scanning, no conditions that would prevent safe and comfortable MR scanning (e.g., implanted medical devices, claustrophobia). Additionally, individuals with ASD were not included if they had a change in their psychotropic medication within the past 6 months. NT individuals with a personal or family history of autism were not included. Two NT participants were taking prescribed antidepressants; excluding these two individuals did not qualitatively affect our results.

Participants who did not achieve criterion performance on catch trials in our behavioral task (1 with ASD, 2 NTs; see Data analysis and statistics section below) were excluded from all analyses (behavioral data, functional MRI, and MR spectroscopy). After exclusion, our final study sample consisted of 28 participants with ASD and 35 NT participants. Demographic information from excluded participants are not included in Table 1. One participant with ASD was excluded from fMRI and MRS analyses (but not behavioral data) due to excessive head motion in the scanner. A summary of missing and excluded data is provided in Supplemental Table 1.

### Visual display and stimuli

Our experimental apparatuses and stimuli have been described previously^1, 71^. Visual experiments were conducted using three different display devices: (1) a ViewSonic PF790 CRT monitor (120 Hz) and Bits# stimulus processor (Cambridge Research Systems, Kent, UK) were used for all psychophysical experiments outside of the scanner. These stimuli were created and displayed in MATLAB (MathWorks, Natick, MA) and PsychToolbox 3^75-77^. Visual stimuli during our fMRI experiments were presented using either (2) an Epson Powerlite 7250 or (3) an Eiki LCXL100A projector (following an equipment failure; both at 60 Hz). These stimuli were created in MATLAB and presented using Presentation software (Neurobehavioral Systems, Berkeley, CA). Viewing distance for all experiments was 66 cm, and luminance was linearized using a PR650 spectrophotometer (Photo Research, Chatsworth, CA).

Stimuli were sinusoidally modulated luminance gratings presented on a mean luminance background (Figure 1A & B). In our psychophysical paradigm outside of the scanner, vertically oriented gratings drifted either left or right (drift rate = 4 cycles/s) within a circular aperture, which was blurred with a Gaussian envelope (*SD* = 0.21°). We used three different stimulus sizes: 0.84, 1.7, and 10° in diameter. The Michelson contrast of the gratings was either 3% (low) or 98% (high), and the spatial frequency was 1.2 cycles/°. Stimuli in the fMRI experiment differed from those in psychophysics as follows: diameter = 2 or 12°, spatial frequency = 1 cycle/°, Gaussian envelope *SD* = 0.25°.

### Experimental protocol

#### Psychophysics

Our psychophysical paradigm, designed to measure spatial suppression, follows the methods of Foss-Feig and colleagues^3^ and has recently been described^1, 71^. In this task, participants were asked to discriminate the direction of motion (left or right) of a briefly presented drifting grating (Figure 1C). Trials began with a shrinking circle fixation mark (850 ms) at the center of the screen, followed by a vertical grating, and then a response period (no time limit). Grating duration was adjusted across trials (range 6.7 to 333 ms) according to an adaptive (Psi) staircase procedure^78^ implemented within the Palamedes toolbox^79^. Correct responses tended to yield shorter durations on subsequent trials. In this way, grating duration was adjusted in order to find the briefest presentation for which the participant would perform with 80% accuracy. Each staircase was composed of 30 trials. Six independent staircases (3 sizes x 2 contrasts) were included in each run and were randomly interleaved across trials. Each run also included 10 catch trials (large, high contrast gratings, 333 ms duration). These low-difficulty trials were intended to measure off-task performance. A total of 4 runs were included in each experimental session, which began with a set of examples and practice trials. Total task duration was approximately 30 min. Data were not obtained in the smallest stimulus size conditions for 5 participants in the ASD group and 5 NTs (for a summary of missing and excluded data, see Supplemental Table 1).

#### Functional MRI

Our fMRI paradigm was designed to measure spatial suppression and has also been described previously^1^. In this task, smaller (2° diameter) and larger (12°) drifting gratings were presented at the center of the screen in alternating 10 s blocks. Grating duration was 400 ms; inter-stimulus interval (ISI) was 225 ms. There were 16 gratings in each block, which drifted in 1 of 8 possible directions (order randomized and counterbalanced). A single fMRI scanning run (4.2 min long) included a total of 25 blocks (13 smaller, 12 larger). Stimulus contrast was either 3% or 98% in separate runs. No baseline or rest blocks were included; stimuli appeared within the central 2° in all blocks. Previous studies^80, 81^ have used this type of alternating block design to measure surround suppression in early visual cortex using fMRI. We chose this paradigm to measure fMRI suppression in ASD for its simplicity and because it allowed us to easily exclude particular blocks from analysis (see below). Each participant completed 2-4 runs at each contrast level across 1 or 2 scanning sessions (some participants chose to end the experiment early, e.g., due to fatigue).

During fMRI, participants performed a color-shape conjunction task at fixation, responding to a green circle in a series of briefly presented colored shapes (e.g., blue square, green square, purple circle). Shapes (0.5° diameter) were presented at the center of the screen (i.e., on top of the gratings) for 200 ms every 1333 ms. Participants responded to the presentation of a green circle by pressing a button on an MR-compatible response pad (Current Designs, Philadelphia, PA). This task encouraged participants to keep their eyes and spatial attention fixed at the center of the screen. We also sought to emphasize bottom-up stimulus processing of the grating stimuli by diverting attention toward the colored shapes in the fixation task.

Our fMRI experiment also included two functional localizer scans, which were used to identify regions of interest (ROIs). The first localizer was designed to identify the motion-selective brain area known as human MT complex (hMT+; Supplemental Figure 1A). We refer to this area as hMT+ to indicate that we did not attempt to differentiate regions MT and MST^82^. This localizer consisted of alternating 10 s blocks of drifting and static gratings (2° diameter, 15% contrast; Supplemental Figure 1B). There were 25 blocks total (13 static, 12 drifting). Grating duration was 400 ms with a 225 ms ISI. The second localizer scan was used to identify voxels with retinotopic selectivity for the central 2°. Using a differential localizer approach^68, 83^ allowed us to identify voxels that responded more strongly to stimuli in the center vs. surrounding portion of the screen. This scan consisted of alternating 10 s blocks of phase-reversing checkerboards (8 Hz; 100% contrast) within the central 2°, or within an annular region from 2-12° eccentricity (Supplemental Figure 1C & D). There were 16 blocks in the second localizer scan (8 center, 8 annulus). Rest blocks were not included during either localizer. Participants performed the same fixation task as in the main fMRI experiment during both localizers. One run of each localizer type was included in each scanning session.

Prior to MR scanning, participants completed a 30 min mock scanning session in which they were introduced to the scanner sounds and practiced lying still in a simulated scanner environment. Participants wore a 3D Guidance trakSTAR motion sensor (Ascension Technology Corp., Shelburne, VT) mounted on a headband. Visual feedback on head motion was given using MoTrak 1.0.3.4 software (Psychology Software Tools, Inc., Sharpsburg, PA). Participants were instructed to keep a small dot representing their head position within a bullseye target region. The mock scanning session also included a practice session for the fMRI fixation task, with examples of the colored shape stimuli, as well as the gratings and checkerboards from each of the fMRI scans.

During the fMRI experiment, we examined fMRI data for head motion as soon as they were acquired. Immediately after each run, data were transferred off the scanner and motion correction was performed using BrainVoyager (see below). Runs in which substantial and sudden head movements (e.g., > 2 mm across 2-4 s) were detected were excluded and repeated within the same scanning session. In such cases, participants were given feedback and coached to remain as still as possible during the subsequent run.

MR data were acquired on a Philips 3 tesla scanner. Each scanning session began with a T_1_-weighted anatomical scan (1 mm isotropic resolution), followed by whole-brain gradient echo fMRI (3 mm isotropic resolution, 30 oblique-axial slices with a 0.5 mm gap, 2 s repetition time [TR], 25 ms echo time [TE], 79° flip angle, anterior-posterior phase encoding direction). A single run with the opposite phase encoding direction (posterior-anterior; 3 TRs) was also acquired during each scanning session, to facilitate geometric distortion compensation.

#### MR spectroscopy

We conducted a ^1^H MR spectroscopy experiment designed to measure GABA+ within particular brain regions. We refer to this metric as GABA+ to indicate that it reflects GABA plus co-edited macromolecules, which are not differentiated by this approach^38^. Our methods were described in our recent publications^1, 71^. Briefly, we used a MEGA-PRESS sequence^84^ to obtain edited MRS data within a 3 cm isotropic voxel (320 averages, 2 s TR, 68 ms TE, 2048 spectral data points, 2 kHz spectral width, 1.4 kHz refocusing pulse, VAPOR water suppression). 14 ms editing pulses were applied at 1.9 ppm (‘on’) or 7.5 ppm (‘off’) during alternating acquisitions within a 16-step phase cycle. The duration of a single MRS run was approximately 11 min. An in-session anatomical scan (as above) was acquired prior to the MRS runs.

We acquired MRS data in two regions of visual cortex: (1) the region of the lateral occipital lobe surrounding area hMT+ (Figure 3A & B; acquired bilaterally in separate runs), and (2) a region of early visual cortex (EVC; Supplemental Figure 2A & B) in the medial occipital lobe aligned parallel and positioned dorsal to the cerebellar tentorium. The hMT+ MRS voxel was placed based on an in-session functional localizer fMRI scan designed to identify area hMT+ (as above, but with duration = 195 s, TR = 3 s, resolution = 3 × 3 × 5 mm, 14 slices with 0.5 mm gap). The hMT+ region was identified online at the scanner using Philips iViewBOLD to identify voxels in the lateral occipital lobe that responded significantly more strongly to moving vs. static gratings (*t* ≥ 3.0). Functional localizer data during the MRS experiment were acquired prior to the anatomical scan and subsequent MRS runs, in order to minimize the effect of frequency drift during MRS caused by gradient heating during fMRI. EVC MRS voxels were positioned according to anatomical landmarks. In order to maximize compliance during MRS, participants watched a theatrical film of their choice to reduce boredom and fatigue. Although we assume that GABA+ values measured with MEGA-PRESS at 3T are relatively stable within individuals^85-87^, differences in visual stimulation between participants may be a source of unaccounted variance in our MRS data.

#### Computational modeling

We applied the normalization model developed by Reynolds and Heeger^24^ to describe motion duration threshold data, as in our previous work^1^. A model diagram is provided in Figure 5. The model can be summarized by the equation:

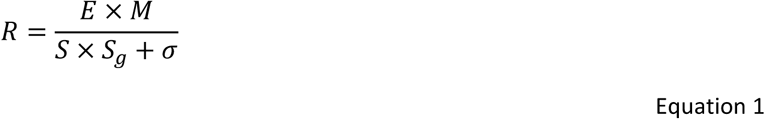

where *R* is the predicted model response (in arbitrary units), *E* is the feed-forward excitatory drive (a function of the visual stimulus strength), *M* is the top-down gain modulation field (a parameter that scales *E* and reflects how stimulus processing is modulated by top-down factors). *S* is the suppressive drive (which depends on the excitatory drive [*E*], but is spatially broader, representing the contribution of a broad ‘normalization pool’), *S*_*g*_ is the suppressive gain (a scaling factor for the suppressive drive, representing the strength of normalization), and *σ* is the semi-saturation constant (a small number that prevents the function from being undefined when the value of *S* is zero). We note that the spatial selectivity of *E* is determined by a Gaussian function with a width parameter *x*_*w_e*_ (Supplemental Equation 2); we refer to this parameter as the width of the excitatory spatial filter (SF; akin to a neural receptive field). This model differs from that in our previous work^*1*^ *in two important ways: 1) the inclusion of the suppressive gain parameter S*_*g*_, and 2) the addition of the top-down modulation parameter *M*. Please see the Supplemental methods for full model details, and Supplemental Table 2 for all parameter values.

To predict duration thresholds from model responses, we assumed an inverse relationship between response magnitude and duration thresholds, such that:

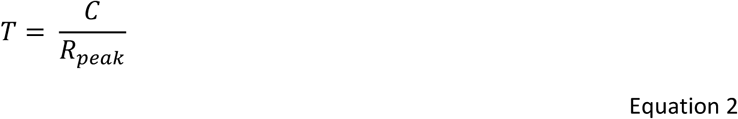

where *T* is the predicted model threshold (in arbitrary units), *C* is the criterion response value needed to reach a perceptual judgment (i.e., left-vs. rightward motion direction)^88^, and *R*_*peak*_ is the peak region of the predicted model response from Equation 1. This inverse relationship between threshold and response is consistent with previous models of motion duration thresholds^4, 46, 89^, and with electrophysiological data from nonhuman primates recorded in area MT during a comparable motion discrimination task^32^.

We compared three different model variants (Figure 6; Supplemental Table 2) to determine which might best match our observation of weaker spatial suppression during motion discrimination in people with ASD vs. NTs (Figure 2A-C). First, we considered the effect of weaker normalization (i.e., a 25% reduction in the suppressive gain term *S*_*g*_) on duration thresholds predicted by the model (see Figure 5, orange box; Figure 6A-C). Using the normalization model, Rosenberg and colleagues^2^ proposed that such a reduction in suppressive gain might account for lower motion duration thresholds in ASD, as reported by Foss-Feig and colleagues^3^. Next, we examined the effect of larger excitatory spatial filters in the model (25% increase in *x*_*w_e*_, the width of the Gaussian function that determines the spatial selectivity of *E*; Figure 5, magenta box; Figure 6D-F). Schauder and colleagues^4^ found that larger spatial filters were able to explain their observation of higher duration thresholds in people with ASD vs. NTs. Finally, we examined whether changing the width of the top-down modulation parameter *M* might affect spatial suppression as predicted by the normalization model (Figure 5, cyan box; Figure 6G-I). In particular, we used a narrower width for the Gaussian parameter that determines the spatial selectivity of *M* in our model (6 vs. 14 arbitrary units). This is consistent with the idea of narrower top-down processing (e.g., spatial attention) during visual perception in ASD, as suggested by previous experimental findings^43^. These three model variants were compared for a *qualitative* match to the pattern of motion duration thresholds we observed in people with ASD vs. NTs (Figure 2A-C). Although other model variants have been successfully used to describe motion duration threshold data (e.g., divisive models with different contrast sensitivity for excitation and suppression^4, 46^), normalization models such as the one applied here have also proven effective^1, 2^. We chose to use a normalization model here in order to directly test hypotheses presented in earlier theoretical work (e.g., the idea that ASD is associated with weaker normalization^2^) against our current experimental data.

#### Clinical measures

Clinical and cognitive assessments were conducted by clinicians with expertise in the evaluation of individuals with neurodevelopmental disorders, and who achieved research reliability on the ADOS-2 and ADI-R; autism diagnoses were confirmed by a trained doctorate-level clinical psychologist using all available information. Overall autism symptom severity was estimated using the ADOS-2 total comparison score^40^. To examine sensory sensitivity and sensory avoidance we used the corresponding domains from the Sensory Profile^41, 42^. Because these two subscales were highly correlated in our sample (*r*_60_ = 0.82, *p* = 3 × 10^-16^), we summed them in order to treat them as a single, combined measure of sensory dysfunction.

### Data analysis and statistics

Psychophysical data were analyzed in MATLAB using the Palamedes toolbox^79^. Duration thresholds for motion direction discrimination were calculated by fitting a Weibull function to the data from each individual staircase. Guess rate and lapse rate were fixed at 50% and 4%, respectively. Thresholds were calculated from the fit psychometric function as the duration value where the participant performed with 80% accuracy. Threshold values less than 0 or above 500 ms were excluded; a total of 3 thresholds were excluded in this way across all data sets. Catch trials accuracy was assessed separately from the staircase data, to examine off-task performance. Participants with less than 80% accuracy across all 40 catch trials were excluded from all further analyses, including fMRI and MRS. One participant with ASD and 2 NTs were excluded in this manner (for a summary of missing and excluded data, see Supplemental Table 1).

To quantify the effect of increasing stimulus size on motion discrimination performance (i.e., suppression), size indices were calculated using an established method^1, 3^, according to Equation 3:

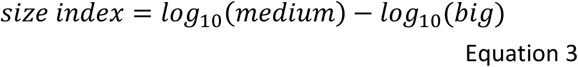

where *medium* indicates the threshold in the medium size condition (1.7° diameter) for a particular contrast level (either 98% or 3%), and *big* indicates the threshold for the big size condition (10° diameter) for the same contrast. More-negative size indices indicate stronger suppression (i.e., a bigger increase in motion duration thresholds with increasing stimulus size).

Functional MRI data were processed in BrainVoyager (Brain Innovation, Maastricht, Netherlands). This included motion correction, distortion compensation, high-pass filtering (> 2 cycles / run), and alignment to the in-session anatomy. Spatial smoothing and normalization to a canonical template were not performed; all analyses were based on within-subject ROIs. ROIs were defined in each hemisphere in the space of the functional data using a standard correlational analysis^1, 68^, taking the top 20 most-significant voxels with an initial threshold of *p* < 0.05 (Bonferroni corrected for multiple comparisons). In a few cases, there were not 20 voxels that satisfied this threshold, thus the threshold was relaxed to include 20 contiguous voxels. All ROIs satisfied a minimum threshold of *p* < 0.006 (1-tailed, uncorrected). EVC ROIs in 4 participants with ASD and 1 NT did not satisfy this criterion; these participants were excluded from EVC fMRI analyses (for a summary of missing and excluded data, see Supplemental Table 1).

ROI location was verified through visualization on an inflated cortical white matter surface model. Bilateral ROIs were defined for area hMT+ in the lateral occipital lobe (Figure 3A & B; Supplemental Figure 1A & B) from the motion vs. static functional localizer data, and for EVC near the occipital pole from the center vs. surround functional localizer data. ROIs in hMT+ were further refined by finding voxels within hMT+ that additionally showed significant retinotopic selectivity for the central 2° in the center vs. surround functional localizer (1-tailed *p* < 0.05; Supplemental Figure 1C & D). Thus, the hMT+ ROIs were identified from the intersection of voxels in the lateral occipital lobe showing selectivity for motion > static, and center > surround. Center-selective regions within hMT+ could not be identified in 3 participants with ASD and 10 NTs, who were thus excluded from hMT+ fMRI analyses (for a summary of missing and excluded data, see Supplemental Table 1).

We extracted fMRI data from each ROI for further analyses in MATLAB using BVQXTools. We examined how fMRI signals within each ROI changed when the stimulus size increased. Within each experimental condition (i.e., high and low stimulus contrast), ROI time series data were divided into epochs spanning 4 s before to 12 s after the stimuli changed from smaller to larger (event-related time = 0 s). Response baseline was calculated by averaging the signal from 0 to 4 s before the size change across all epochs. We converted the data to percent signal change by subtracting and then dividing by the baseline value, and then multiplying by 100. To compute an average response time course in each condition for each participant, we took the mean signal for each time point across all epochs, hemispheres and fMRI runs. The magnitude of the fMRI response to the increase in stimulus size was calculated as the average signal from 8 to 12 s after the size increase (the time period when suppression was maximal).

MRS data were analyzed in the Gannet 2.0 Toolbox^90^ within MATLAB. Data were processed using the toolbox-standard approach, including automated frequency and phase correction, artifact rejection (frequency correction > 3 *SD* above the mean), and 3 Hz exponential line broadening. To calculate the concentration of GABA+, we fit a Gaussian to the peak in the MEGA-PRESS spectrum at 3 ppm (Supplemental Figure 5). We refer to this value as GABA+ to indicate that it reflects GABA plus co-edited macromolecules which are not differentiated by this method. Likewise, the Glx peak at 3.75 ppm was fit with a double Gaussian. The area under the fit curve served as a measure of the metabolite level. GABA+ and Glx were each scaled relative to water; the unsuppressed water peak was fit with a mixed Gaussian-Lorentzian. Tissue correction was performed for GABA+ based on the proportion of gray matter, white matter, and CSF within each MRS voxel, assuming twice the concentration of GABA+ in gray vs. white matter, using an established method^91^. Tissue fraction values within each MRS voxel were obtained by segmenting the T_1_ anatomical scan using SPM8^92^. There is currently no standard method for tissue correction for Glx. Instead, we performed a series of control analyses to explore the contributions of different tissue types and the measured water reference signal to our Glx measurements (see Supplemental information). Concentrations for GABA+ and Glx are reported in institutional units (i.u.). We collected all psychophysical, fMRI, and MRS data within a 2-week time period for each participant; previous studies suggest that GABA+ values are fairly stable over this time period^85-87^. One participant with ASD was excluded from MRS analyses due to excessive head motion, as evidenced by large water frequency shifts across time (for a summary of missing and excluded data, see Supplemental Table 1).

Statistical analyses were performed in MATLAB. Normality and homogeneity of variance were assessed by visual inspection of the data. Group differences were assessed using mixed repeated measures analyses of variance (ANOVAs). Participants were modeled as a random effect and nested within groups. Stimulus size was modeled as a continuous variable. Reported *r*-values are Pearson’s correlation coefficients; associated *p*-values were determined using non-parametric 2-tailed permutation tests, unless otherwise noted. Here, we randomly shuffled the data being correlated across participants (without replacement) in each of 10,000 iterations; *p*-values were calculated as the proportion of shuffled samples with *r*-values more extreme than the observed *r*-value. Bonferroni correction was used to adjust *p*-values for multiple comparisons.

## Acknowledgments

We thank Brenna Boyd, Judy Han, Ly Nguyen, Heena Panjwani, Micah Pepper, Meaghan Thompson, Anne Wolken, and the UW Diagnostic Imaging Center for help with recruitment and/or data collection. We thank Geoffrey M. Boynton for providing the MATLAB functions for the computational model.

This work was supported by funding from the National Institutes of Health (F32 EY025121 to MPS, R01 MH106520 to SOM, T32 EY00703). This work applies tools developed under NIH grants R01 MH098228, R01 EB016089, and P41 EB015909; RAEE also receives support from these grants.

## Supplemental information

### Supplemental results

#### Additional hMT+ MRS analyses

As MRS data were acquired in both left and right hMT+ in separate scans, we compared data between hemispheres to examine potential lateralized differences in metabolite concentrations, or interactions between participant group and hemisphere. We found no difference in GABA+ levels between left and right hMT+ across groups (GABA+: left mean = 2.54 i.u., *SD* = 0.19; right mean = 2.54 i.u., *SD* = 0.23; main effect of hemisphere, *F*_1, 60_ = 9 × 10^-5^, *p* = 1.0). In contrast, Glx levels were higher in right as compared to left hMT+ (Glx: left mean = 1.18 i.u., *SD* = 0.19, right mean = 1.44 i.u., *SD* = 0.14; main effect of hemisphere, *F*_1, 60_ = 107, *p* = 6 × 10^-15^). There were no significant interactions between group & hemisphere for either metabolite (*F*_1, 60_ < 1.42, *p*-values > 0.2).

**Supplemental Figure 1.**
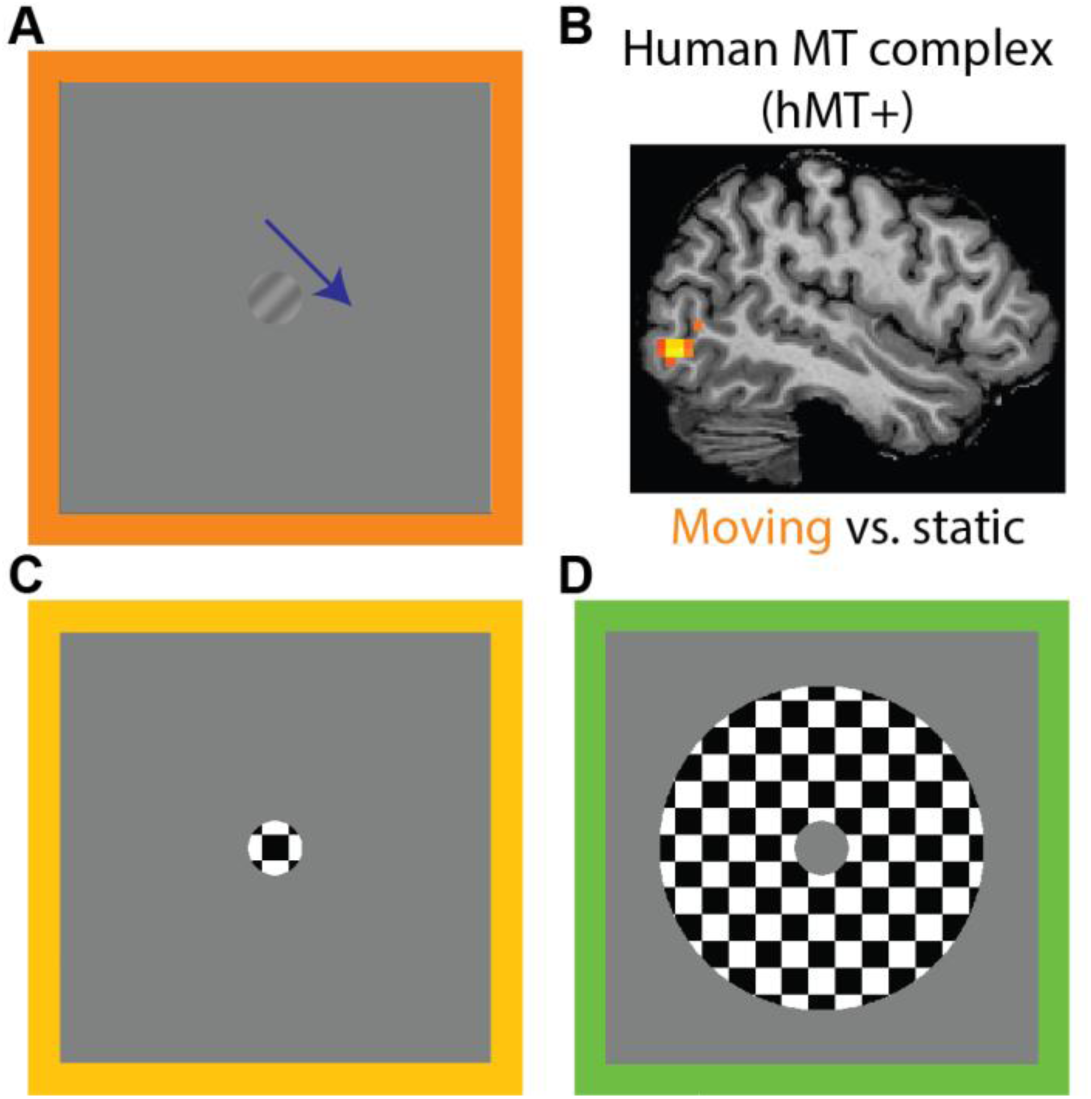
Functional localizers. **A)** Functional localizer scan #1 included blocks of moving vs. static gratings. **B)** Human MT complex (hMT+) was identified in the lateral occipital lobe using a correlational analysis (motion > static). Functional localizer scan #2 included blocks of flickering checkerboards in the central 2° **(C)**, vs. surrounding 12° **(D)**. Foveal hMT+ ROIs were defined by (motion > static) & (center > surround).

Because our GABA+ values were corrected for tissue fraction, while our Glx values were not (see Methods), we asked whether differences in gray matter, white matter, and CSF fractions within the voxel might have contributed to the observed difference in Glx levels between left and right hMT+. We found that the proportion of CSF was indeed higher within left vs. right hMT+ voxels (left mean = 3.79%, *SD* = 1.99%; right mean = 3.30%, *SD* = 1.54%; *F*_1, 60_ = 8.77, *p* = 0.004). There were no significant differences in gray or white matter content between hemispheres (GM: left mean = 44.5%, *SD* = 3.67%; right mean = 44.3%, *SD* = 3.24%; *F*_1, 60_ = 0.14, *p* = 0.7; WM: left mean = 51.7%, *SD* = 4.22%; right mean = 52.4%, *SD* = 3.51%; *F*_1, 60_ = 2.64, *p* = 0.11). Differences in CSF were reflected in the water reference; the measured water signal was also higher in left vs. right hMT+ (left mean = 30.5 i.u., *SD* = 5.33, right mean = 28.9 i.u., *SD* = 5.46; *F*_1, 60_ = 27.4, *p* = 2 × 10^-6^). However, there were no significant differences between participants with ASD and NTs in gray matter, white matter, CSF, or water levels (*F*_1, 60_ < 3.54, uncorrected *p*-values > 0.065), and no significant interactions between group and hemisphere (*F*_1, 60_ < 2.25, uncorrected *p*-values > 0.139). Thus, although differences in Glx values from left vs. right hMT+ depended on CSF fraction and the associated water reference signal, we did not find evidence to suggest that these effects were associated with any differences in MRS data between groups.

To further examine whether our MRS results in hMT+ were affected by scaling our metabolite values (GABA+ and Glx) relative to water, we repeated our analyses using values instead scaled relative to creatine. We found comparable results for the creatine scaled data; there were no group differences in GABA+ (*F*_1, 60_ = 1.42, *p* = 0.2) or Glx (*F*_1, 60_ = 0.84, *p* = 0.4), and no correlations with suppression metrics (size indices or fMRI suppression, averaged across contrast conditions to minimize multiple comparisons, |*r*_47-60_| < 0.16, uncorrected *p*-values > 0.2; data not shown). Thus, we found no evidence to suggest that GABA+ or Glx levels in hMT+, as measured by MRS, differed between participants with ASD and NT controls.

#### MRS in EVC

Additional MRS data were acquired in a mid-occipital voxel within early visual cortex (EVC; Supplemental Figure 2A & B). We first examined whether metabolite values (GABA+ or Glx) in EVC differed between groups. No significant group differences were observed (main effect of group, GABA+: *F*_1, 60_ = 1.11, *p* = 0.3; Glx: *F*_1, 60_ = 0.55, *p* = 0.5; Supplemental Figure 2C & D). We also examined correlations between GABA or Glx in EVC and size indices from our psychophysical task, none of which were significant (correlations examined both within and across participant groups, size indices averaged across contrast conditions to minimize multiple comparisons, |*r*_25-60_| < 0.24, uncorrected *p*-values > 0.15; data not shown). These results do not suggest that differences in the strength of spatial suppression during motion discrimination, either between individuals or between participants with ASD and NTs, may be explained by differences in GABA+ or Glx levels within EVC as measured by MRS.

#### Computational modeling

To reconcile our current modeling work with previous studies^2, 4^, we asked whether spatially narrower top-down modulation within the normalization model might be sufficient to account for previous findings of lower^3^ or higher^4^ motion duration thresholds in ASD. Indeed, we found that using an overall narrower set of top-down modulation parameters (2 vs. 6 arbitrary units) yielded model predictions (Supplemental Figure 3A-C) that showed a good qualitative match to the results of Foss-Feig and colleagues^3^. Similar to our model predictions, they observed smaller duration thresholds across stimulus sizes for high contrast gratings, and for large, low contrast stimuli. Additionally, we found a different pattern of results using an even narrower top-down modulation field (1 vs. 6 arbitrary units). Model duration thresholds differed less at high contrast, and were actually higher at low contrast for very-narrow vs. broader top-down modulation in our model (Supplemental Figure 3D-F). This occurs because the top-down modulation field is actually narrower than the response region that is ‘read out’ (i.e., averaged when computing the predicted threshold; *r*_*w*_; Supplemental Equation 5). The fact that narrower top-down processing can actually yield higher predicted duration thresholds within the model framework may therefore help to reconcile disparate observations of lower^3^ and higher^4^ motion duration thresholds among people with ASD vs. NTs.

**Supplemental Figure 2.**
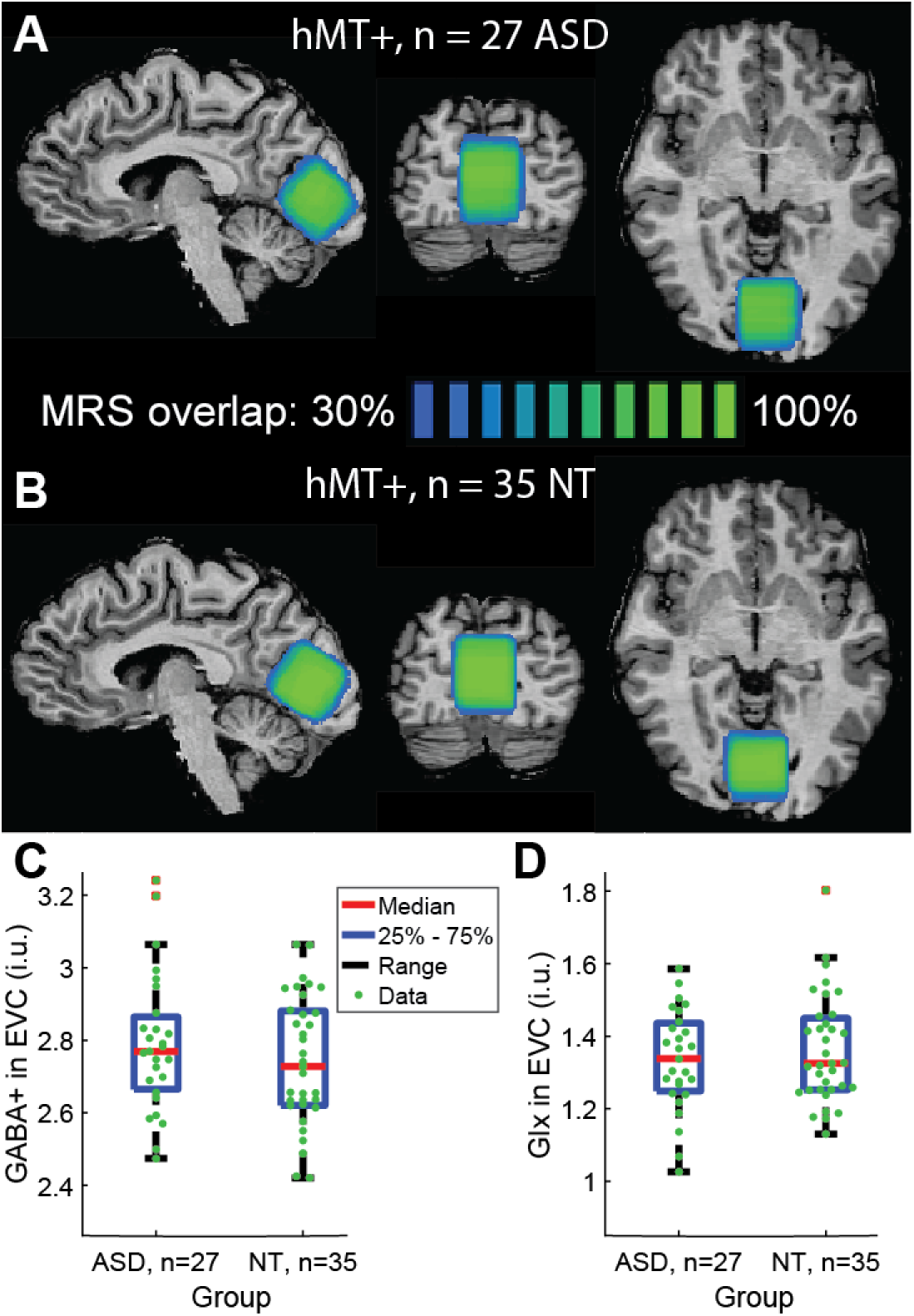
MRS results in EVC. **A)** Average voxel positions for EVC in Talairach space for n = 27 participants with ASD. Blue-green color indicates percent overlap across individuals. **B)** Same, but for n = 35 NT participants. **C)** GABA+ concentrations in EVC. Values are in institutional units (i.u.). **D)** Same, but for Glx.

#### Control analyses

Previous studies have suggested that age^44-46^, biological sex^49^, and IQ^49-51^ may each influence motion discrimination thresholds. None of these demographic factors differed significantly between our groups of participants with ASD and NTs (Table 1). Nevertheless, to further control for these factors, we conducted post-hoc analyses to examine group differences in duration thresholds, size indices, and fMRI suppression, with age, sex, and non-verbal IQ included as covariates. In each case, the results reported in the main text were recapitulated in the post-hoc analyses; we saw a significant interaction between group and size in our analysis of motion duration thresholds, and significant group differences for size indices and fMRI suppression in hMT+ (linear mixed-effect models, thresholds: interaction between group and size, parameter estimate [SE]: 3.77 [0.59], *t*_1420_ = 6.40, *p* = 2 × 10^-10^; size indices: main effect of group, parameter estimate [SE]: 0.152 [0.042], *t*_497_ = 3.60, *p* = 4 × 10^-4^; fMRI suppression: main effect of group, parameter estimate [SE]: 0.176 [0.061], *t*_93_ = 2.91, *p* = 0.005). These post-hoc results suggest that our observations of weaker suppression during motion discrimination and in the fMRI response within hMT+ among participants with ASD vs. NTs are not explained by subtle differences between groups in demographic factors such as age, sex, or IQ.

We also considered whether our fMRI results showing weaker suppression in hMT+ in participants with ASD might be explained by differences in head motion or a lack of engagement in the color-shape conjunction task (performed at fixation). To this end, we excluded fMRI blocks (10 s) with excessive head motion, and scans (4 min) in which there was poor fixation task performance (see Supplemental methods for details). Data from 2 ASD participants and 1 NT were completely excluded, as there was not a sufficient amount of data for analysis following the above exclusion. Despite the reduction in sample size, the results of this secondary analysis were qualitatively the same as those reported for the full data set (Figure 2D-F); in response to larger drifting gratings, suppression of the fMRI signal in hMT+ was weaker in participants with ASD (Supplemental Figure 4; ANOVA, main effect of group: *F*_1, 44_ = 4.10, *p* = 0.049). This suggests that differences in head motion or fixation task performance between groups may not account for weaker fMRI suppression within hMT+ in ASD.

**Supplemental Figure 3.**
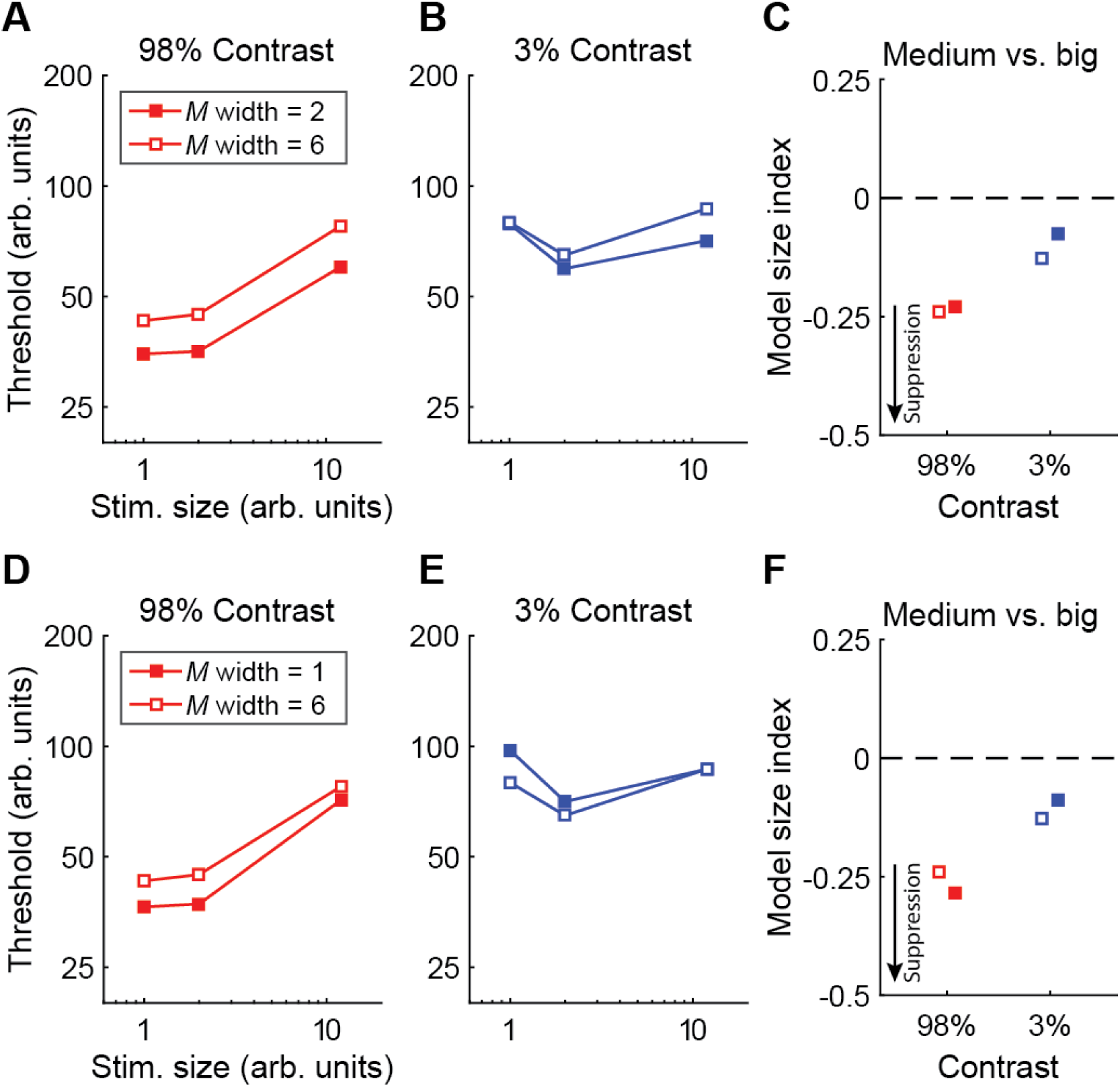
Supplemental model results. **A-C)** A narrower set of top-down modulation field parameters (the spatial width of the parameter *M* from Supplemental Equation 1, abbreviated *M* width; 2 vs. 6 arbitrary units) yields model predictions that are reasonably well matched to the observations of Foss-Feig and colleagues^3^; duration thresholds are lower across sizes at high contrast, and for large, low contrast stimuli. **D-F)** Further reducing the width of the narrower top-down modulation (to 1 arbitrary unit) reveals a different, mixed pattern; duration thresholds are lower for high contrast, but higher for low contrast stimuli.

Finally, we examined whether differences in eye movements between groups might account for weaker spatial suppression in our participants with ASD. We conducted a series of analyses to examine relationships between suppression metrics from our behavioral and fMRI experiments, and eye tracking data collected simultaneously. We used 3 eye tracking metrics: mean distance from center (a measure of gaze accuracy in space), the standard deviation of fixation distance from center (a spatial measure of gaze variability), and mean fixation duration (a measure of fixation stability across time). Eye tracking data were drift corrected post-hoc (see Supplemental methods for eye tracking analysis details). To assess whether fixation behavior may have influenced our observation of weaker spatial suppression in ASD, we examined group differences in eye tracking metrics, and correlations between these metrics and psychophysical or fMRI measures of suppression across both ASD and NT participants.

Eye tracking data were obtained during psychophysics for 14 participants with ASD and 25 NTs. We found no evidence of reliable differences in eye tracking metrics during psychophysics between groups (Mann-Whitney tests, *Z*-values < 0.63, uncorrected *p*-values > 0.5, data not shown). Eye tracking metrics were also not correlated with psychophysical suppression measures (SI at 98% or 3% contrast) across participants (|*r*_37_ < 0.28|, uncorrected *p*-values > 0.091, data not shown).

Eye tracking data were obtained during fMRI for 14 participants with ASD and 24 NTs. The drift corrected mean and *SD* of distance from fixation were numerically higher for ASD vs. NT participants, but these differences were not statistically significant (Mann-Whitney tests, *Z*-values < 1.59, uncorrected *p*-values > 0.11, data not shown). There was also no reliable difference in fixation time between groups (*Z* = 0.5, *p* = 0.6). Of the participants for whom eye tracking data were collected during fMRI, foveal MT ROIs were identified for 12 individuals with ASD and 19 NTs. Eye tracking metrics were not significantly correlated with fMRI suppression within hMT+ (at 98% or 3% contrast) across participants (|*r*_29_ < 0.33|, uncorrected *p*-values > 0.071, data not shown). Although the conclusions that may be drawn from our eye tracking data are limited by the small sample size, we do not find evidence to suggest that weaker spatial suppression in ASD may be explained by gross systematic differences in fixation behavior during psychophysical or fMRI task performance.

**Supplemental Figure 4.**
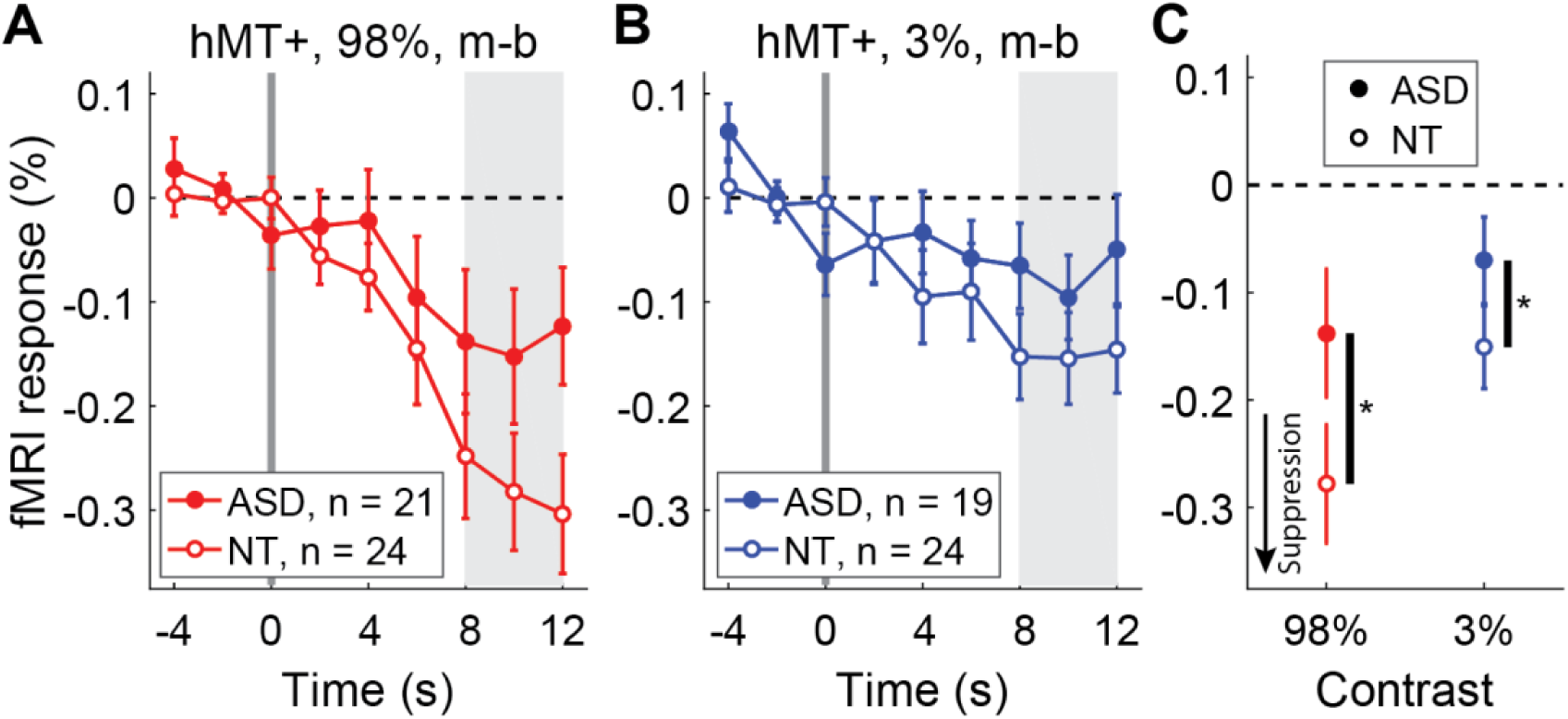
FMRI results with blocks excluded for excess head motion and poor task performance. **A)** As in Figure 2, fMRI responses in foveal Hmt+ to an increase in stimulus size; at time = 0 s, high contrast drifting gratings increased in size from medium (m) to big (b). **B)** The same, but for low contrast gratings. **C)** Average fMRI responses for each group (from shaded regions in **A** & **B**). Error bars are *S.E.M.*

### Supplemental methods

#### Missing or excluded data points

Supplemental Table 1 summarizes the missing or excluded data points from both participant groups across all experiments. Details for the exclusion procedures are provided in the relevant sections of the Methods in the main text.

**Supplemental Table 1.**
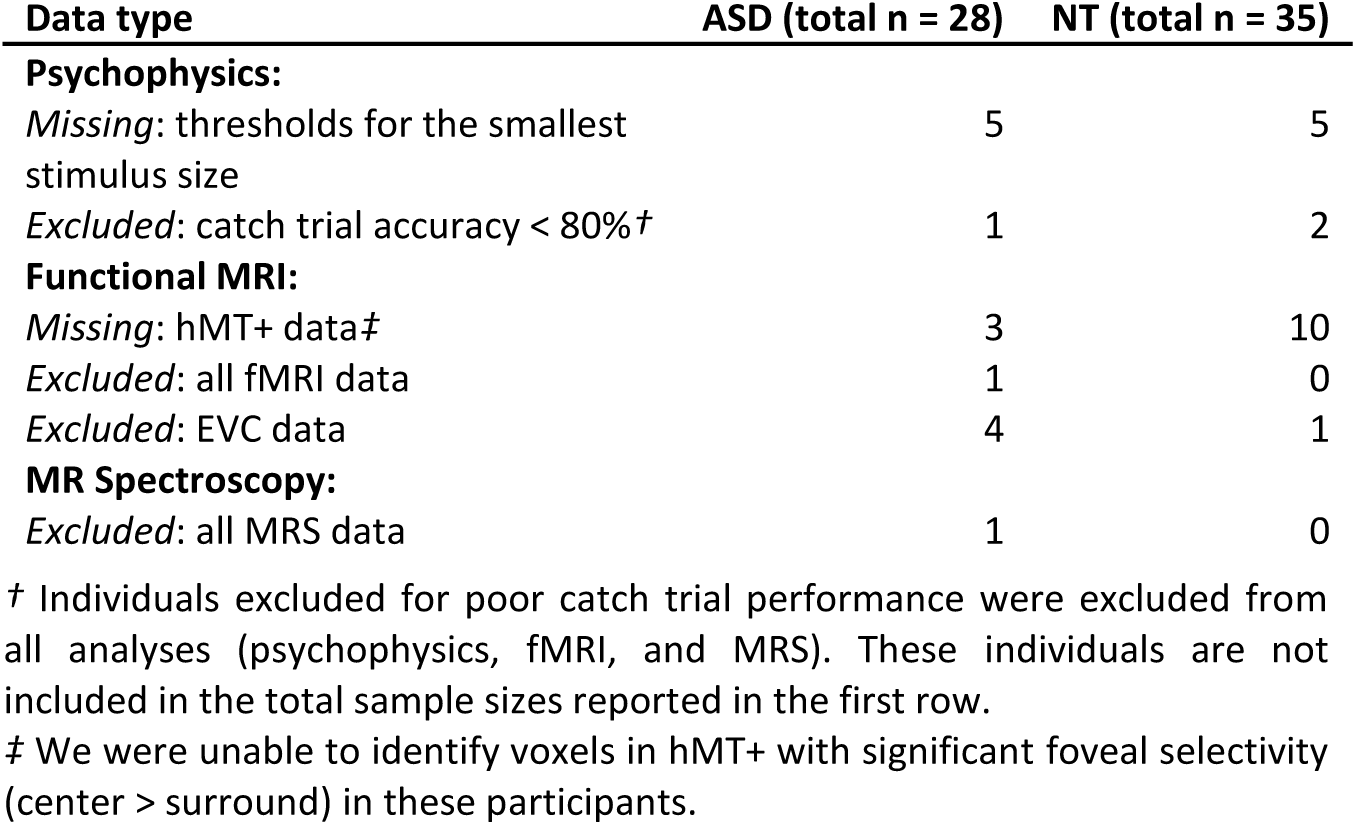
Summary of missing or excluded data. The number of individuals with missing data is reported for each group.

| Data type | ASD (total n = 28) | NT (total n = 35) |
| --- | --- | --- |
| **Psychophysics:** |  |  |
| Missing: thresholds for the smallest stimulus size | 5 | 5 |
| Excluded: catch trial accuracy < 80%<sup>†</sup> | 1 | 2 |
| **Functional MRI:** |  |  |
| Missing: hMT+ data<sup>‡</sup> | 3 | 10 |
| Excluded: all fMRI data | 1 | 0 |
| Excluded: EVC data | 4 | 1 |
| **MR Spectroscopy:** |  |  |
| Excluded: all MRS data | 1 | 0 |

<sup>†</sup> Individuals excluded for poor catch trial performance were excluded from all analyses (psychophysics, fMRI, and MRS). These individuals are not included in the total sample sizes reported in the first row.
<sup>‡</sup> We were unable to identify voxels in hMT+ with significant foveal selectivity (center > surround) in these participants.

#### Computational modeling

Our computational approach is a direct application of the normalization model published by Reynolds and Heeger^24^. We have previously used a similar modeling technique to describe motion duration threshold data across a variety of experimental conditions in NT participants^1^. A summary of the current model approach is provided in the Methods; a thorough description of the method is provided below.

The core of the model, summarized in Equation 1, is reprinted here as:

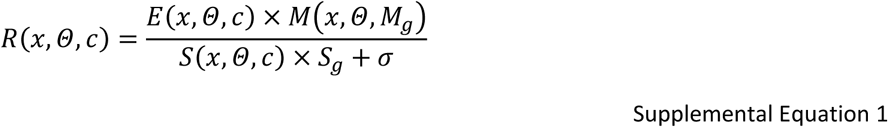

where *R* is the predicted model response as a function of stimulus space (*x*), orientation (*Θ*), and contrast (*c*). The suppressive gain parameter *S*_*g*_ is a scalar that controls the strength of divisive normalization^2^, and *σ* is the semi-saturation constant, which controls the non-linearity of the predicted response as a function of stimulus contrast, as well as preventing the value of *R* from being undefined when *S* equals zero.

The parameters *E, S*, and *M* are each 2-dimensional representations of a population of computational processes, with selectivity across a spatial dimension (*x*) and an orientation dimension (*Θ*). The magnitudes of *E* and *S* are a function of stimulus contrast (*c*). Thus:

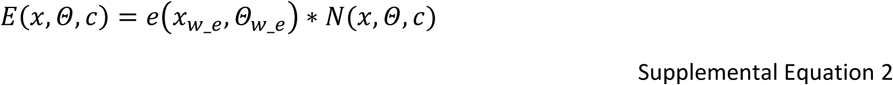

where *N* is the ‘stimulus image,’ which is a population-level representation of the stimulus input for the model, and * denotes convolution. Specifically, *N* is a 2-dimensional Gaussian function whose amplitude is set by *c*. The widths of *N* in the *x* and *Θ* dimensions are determined by the shape of the stimulus being modeled. Likewise, *e* is a 2-dimensional Gaussian function, where the spatial and orientation selectivity for the excitatory drive (*E*) are determined by the width of spatial (*x*_*w_e*_) and orientation (*Θ*_*w_e*_) Gaussian parameters, referred to as excitatory spatial filters and orientation filters, respectively. These filter parameters can be thought of as the model equivalent of the spatial and orientation selectivity of neural receptive fields in visual cortex. The amplitude of *e* is set to 1, such that *c* determines the amplitude of *E*.

As with the stimulus parameter *N*, the top-down gain modulation field parameter *M* is a 2-dimensional Gaussian function: *M*(*x, Θ, M*_*g*_). The center and width of top-down modulation in the spatial (*x*) and orientation (*Θ*) dimensions are determined by the mean and tuning width of *M*(*x*) and *M*(*Θ*), while the amplitude is set by the top-down modulation gain factor *M*_*g*_. The minimum value of *M* is set to 1, such that there is no effect of multiplying *E* x *M* (in the numerator of Supplemental Equation 1) outside of the region where top-down processing is focused (see Figure 5, cyan box).

The suppressive drive *S* is a function of both *E* and *A*, such that:

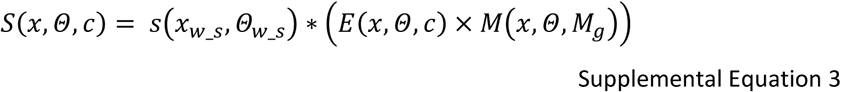

where *s* is another 2-dimensional Gaussian, like *e* in Supplemental Equation 2. Importantly, normalization models^23, 24^ generally assume that the selectivity of *s* (in space; *x*_*w_s*_) is broader than *e*. This produces the characteristic size tuning in the predicted model response, such that increasing stimulus size leads to a peak and then a decrease in the predicted response (see Figure 2F from our recent paper^1^ for a graphical depiction of size tuning from this model).

The predicted motion duration threshold is given by Equation 2, reprinted here:

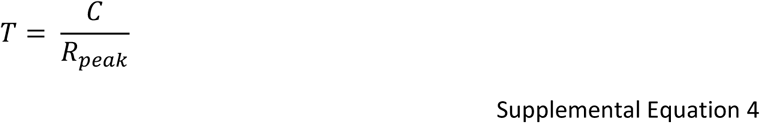

where *T* is the predicted threshold in arbitrary units, and *C* is the criterion response level for the model. The value of *R*_*peak*_ is an average of the region surrounding the peak of the predicted model response *R*, such that:

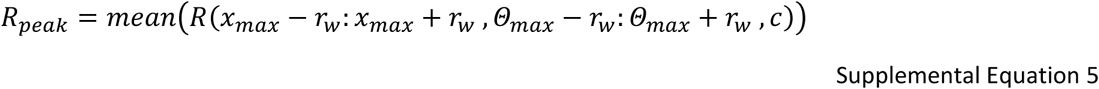

where *R*(*x*_*max*_, *Θ*_*max*_) is the maximum value of *R*, which happens to correspond to the center of the modeled stimulus. The parameter *r*_*w*_ determines the width of the region surrounding the peak in which the average is taken. In essence, using Supplemental Equation 5, we find the mean response in the region of the peak predicted response. While there is empirical support from electrophysiology in non-human primates for the notion that motion duration thresholds depend on the response of neurons whose spatial receptive field is centered on the stimulus^32^, we note that this method differs slightly from our previous work^1^, in which the threshold *T* depended strictly on the maximum of the predicted response *R*. Values for each model parameter across all three model variants (from Figure 6) are provided in Supplemental Table 2, and are adapted from our previous work^1^.

**Supplemental Table 2.** Model parameters. Parameters marked with an asterisk **(*)** varied across the three model versions in Figure 6. Top-down spatial width values in parentheses are from the model variants shown in Supplemental Figure 3. Inf. = infinite.

| Parameter name | Parameter value |  |  |
| --- | --- | --- | --- |
|  | Weak norm. model | Large excitatory SF model | Narrow top-down model |
| Stimulus contrast | 0.03 or 0.98 | 0.03 or 0.98 | 0.03 or 0.98 |
| Stimulus spatial center ($x$, a.u.) | 0 | 0 | 0 |
| Stimulus spatial width (a.u.) | 1, 2, or 12 | 1, 2, or 12 | 1, 2, or 12 |
| Stimulus orientation ($\theta$, °) | -90 | -90 | -90 |
| Stimulus orientation width (°) | 3 | 3 | 3 |
| Excitatory spatial pooling width ($x_{w,e}$, a.u.)* | 4 | 5 vs. 4 | 4 |
| Excitatory orientation pooling width ($\theta_{w,e}$, °) | 25 | 25 | 25 |
| Top-down spatial center ($x$, a.u.) | 0 | 0 | 0 |
| Top-down spatial width (a.u.)* | 14 | 14 | 6 vs. 14<br>(1 or 2 vs. 6) |
| Top-down orientation ($\theta$, °) | -90 | -90 | -90 |
| Top-down orientation width (°) | Inf. | Inf. | Inf. |
| Top-down gain ($M_g$) | 4 | 4 | 4 |
| Suppressive spatial pooling width ($x_{w,s}$, a.u.) | 40 | 40 | 40 |
| Suppressive orientation pooling width ($\theta_{w,s}$, °) | 25 | 25 | 25 |
| Suppressive gain ($S_g$)* | 0.75 vs. 1 | 1 | 1 |
| Semi-saturation constant ($\sigma$, a.u.) | 0.0002 | 0.0002 | 0.0002 |
| Criterion ($C$, a.u.) | 600 | 600 | 600 |
| Response region width ($r_w$, a.u.) | 1 | 1 | 1 |

#### Statistics

We used linear mixed effects modeling in our post-hoc analyses that controlled for age, sex, and non-verbal IQ. With the addition of these 3 factors, we chose to use linear mixed effects models rather than ANOVAs, in order to make our analyses robust against missing factor combinations (i.e., rank deficiency). Linear mixed effects models were fit using a maximum likelihood procedure via the *fitlme.m* function in MATLAB.

#### Functional MRI

To account for possible residual effects of head motion in our fMRI results, we performed control analyses in which we excluded individual fMRI blocks (10 s) in individual participants in which excessive head motion was detected. Although we performed motion correction in BrainVoyager (Brain Innovation, Maastricht, Netherlands) as part of our fMRI preprocessing, such corrections are incomplete at best, and large head movements may still negatively affect data quality even after correction. Our motion correction procedure yielded estimates of translation in x, y, and z dimensions, as well as roll, pitch, and yaw rotations between each pair of subsequent TRs. Framewise displacement^93^ was calculated by taking the sum of the absolute value of the displacement in each of these six dimensions, with rotation converted from degrees to millimeters on the surface of a sphere with a radius of 50 mm. The threshold for excessive head motion was defined as a framewise displacement > 0.9 mm^94^. We excluded all fMRI blocks that contained TRs with any framewise displacement values larger than this threshold. In order to account for the slow time course of the hemodynamic response, framewise displacement data were scrutinized within a time window spanning from 16 s before to 4 s after each 10 s fMRI block.

We also sought to account for possible effects of task disengagement during fMRI by excluding individual fMRI scans in which the participant performed poorly on the fixation task. The task consisted of a color-shape conjunction search, in which the participant responded with a button press to the appearance of a green circle in a series of small, briefly presented colored shapes. Poor task performance was defined based on a threshold of 60% hit rate; we excluded all fMRI scans (4 min) in which performance was below this threshold.

#### Eye tracking

Eye tracking data were acquired during both psychophysics and fMRI using an SR Research (Ottawa, Canada) EyeLink 1000 infrared eye tracking camera system. Data were acquired at a sampling rate of 1000 Hz.

We identified fixation periods using a sliding window analysis along with post-hoc drift correction. Specifically, we used a dispersion-based fixation detection algorithm^95, 96^. Fixations were defined within set periods of at least 100 ms for which the maximum distance of any gaze position measurement within the set compared to the set’s centroid does not exceed a threshold radius of 1°. In this algorithm, subsequent time points were added to a set until this distance threshold was exceeded, at which point a new set was defined. The parameters that defined fixation periods were taken from previous work using this method^96^. Next, post-hoc drift correction was performed by calculating the average gaze position across all fixations within a 10 s period, taking the difference between this average fixation position and the intended fixation position (i.e., the fixation mark at the center of the screen), and subtracting this value from all gaze position measures within the 10 s period. This analysis assumes that participants understood the instructions to maintain fixation at the center of the screen and were attempting to do so throughout the task. We further assumed that systematic differences between measured and instructed gaze position may be accounted for by drift in the eye tracking camera calibration over time (e.g., due to slight postural changes). We believe these assumptions are reasonable given that fixation performance was monitored by study staff, who verified comprehension of the fixation protocol with the participants prior to task initiation. This drift correction method is a subtractive version of an algorithm developed by Vadillo and colleagues^97^. We chose a subtractive method, rather than a multiplicative one (as used in their original paper), as we found the former was better suited to drift correction during a central fixation task, and did not dramatically alter the magnitude of gaze position variability metrics.

We calculated three different eye tracking metrics: mean distance from center (a measure of gaze accuracy in space), the standard deviation of fixation distance from center (a spatial measure of gaze variability), and mean fixation duration (a measure of fixation stability across time). Because we found that these data were not normally distributed, we used Mann-Whitney tests to assess group differences, and Spearman’s Rho to test for correlations.

**Supplemental Figure 5.**
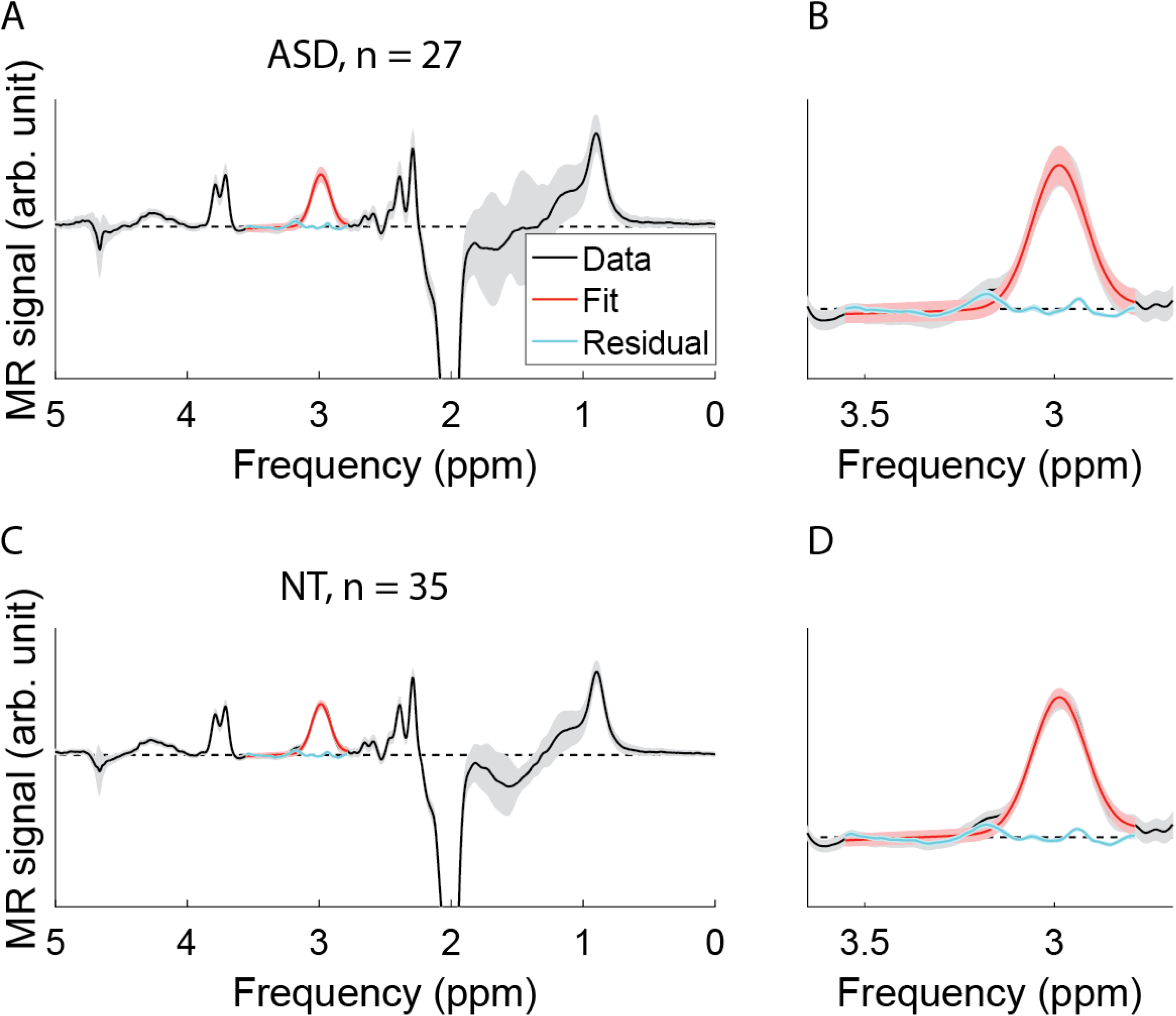
Group average MR spectra. **A)** Average MR spectroscopy data from n = 27 participants with ASD. Data are shown in black, Gaussian fits are shown in red, residuals are shown in blue. Signal intensity is shown in arbitrary units. **B)** Zoomed in section from **A**, to better illustrate fits. Panels **C** & **D** show the same, but for n = 35 NT participants. Shaded areas show 1 *SD*. Axes are scaled equally in panels **A** & **C**, and in panels **B** & **D**.

